# Repair, resilience and asymmetric segregation of damage in the context of replicative ageing: it is a balancing act

**DOI:** 10.1101/446302

**Authors:** Johannes Borgqvist, Niek Welkenhuysen, Marija Cvijovic

**Author notes:** Corresponding Author: Marija Cvijovic, Chalmers University of Technology and the University of Gothenburg, Mathematical Sciences, Chalmers tvärgata 3, SE-412 96 Gothenburg, Sweden.

## Abstract

Accumulation of damaged proteins is a hallmark of ageing, occurring in organisms ranging from bacteria and yeast to mammalian cells. During cell division in *Saccharomyces cerevisiae*, damaged proteins are retained within the mother cell, resulting in a new daughter cell with full replicative potential and an ageing mother with a reduced replicative lifespan (RLS). The cell-specific features determining the lifespan remain elusive. It has been suggested that the RLS is dependent on the ability of the cell to repair and retain pre-existing damage. To deepen the understanding of how these factors influence the life span of individual cells, we developed and experimentally validated a dynamic model of damage accumulation accounting for replicative ageing. The model includes five essential properties: cell growth, damage formation, damage repair, cell division and cell death, represented in a theoretical framework describing the conditions allowing for replicative ageing, starvation, immortality or clonal senescence. We introduce the resilience to damage, which can be interpreted as the difference in volume between an old and a young cell. We show that the capacity to retain damage deteriorates with high age, that asymmetric division allows for retention of damage, and that there is a trade-off between retention and the resilience property. Finally, we derive the maximal degree of asymmetry as a function of resilience, proposing that asymmetric cell division is beneficial with respect to replicative ageing as it increases the RLS of a given organism. The proposed model contributes to a deeper understanding of the ageing process in eukaryotic organisms.

## Introduction

Cell division, growth and death are fundamental features of any living organism. During its life cycle, a cell produces a set of functional components such as proteins or other metabolites, which will ultimately allow the cell to divide, giving rise to a newly born daughter cell. However, due to inherent imperfections in the cellular machinery over time, cells are slowly deteriorating causing essential intracellular functions to perish. At the very end of a cell’s lifespan, age-associated damage builds up consistently impairing the ability of the cell to divide and survive, which eventually culminates in cell death. In environments with a sufficient amount of food, damage will be formed as a consequence of cell growth. This causes the partitioning of damage between the progenitor and progeny during cell division to be an important function.

An asymmetric distribution of cell mass after division constitutes a vital part of ageing in the yeast *Saccharomyces cerevisiae* (*S.cerevisiae*). The number of divisions before cell death is a measure of the age of a single yeast cell, which has been studied substantially by means of experiments [1, 2], and it is called the replicative life span (RLS). The asymmetric division of the budding yeast results in a large ageing mother cell with a finite RLS and a new small daughter cell with full replicative potential [3]. Replicative ageing in yeast is characterised by the accumulation of age-related damage that is selectively retained within the mother cell compartment at each division [4]. This retention of damage is required to rejuvenate daughter cells and thereby maintain viability in populations over time. The described mode of division resembles the division observed in multicellular organisms, characterised by the separation of the soma and the germ cell lines [5]. However, the details behind the interaction of the involved systems and their effects on ageing are still largely unknown. To elucidate the interplay between the various constituents of these processes, theoretical approaches are often implemented in addition to experimental methods.

Hitherto, multiple qualitative mathematical models of the accumulation of damage have been developed [6, 7, 8, 9, 10, 11, 12]. One of the first models showed that even simple unicellular bacteria undergo ageing as a means to cope with damage [6]. This finding suggested that the phenomenon of ageing precedes eukaryotes and has evolved numerous times as it occurs in multiple organisms. A model of damage accumulation showed the importance of asymmetric partitioning of damage through retention in the symmetrically dividing fission yeast *Schizosaccharomyces pombe* (*S.pombe*) and in the asymmetric budding yeast *S.cerevisiae* as it increases the fitness of the population at both low and high damage propagation rates [7]. A model of ageing in the bacteria *Escherichia coli* (*E.coli*) showed that under the examined conditions the cell lineage was immortal due to its low rate of formation of damage [8]. A follow-up model quantified the importance of a deterministic (as opposed to a stochastic) asymmetric distribution of damage upon cell division [11], also further concluding that asymmetric partitioning of damage increases the overall fitness of the population. An additional study in yeast proposed that the ratio between maintenance and growth is critical when determining the benefits of symmetric and asymmetric division [9]. Furthermore, damage repair has been identified as a better strategy in terms of coping with damage compared to asymmetric segregation [10]. Most recently, it was demonstrated that asymmetric segregation of damage in *E.coli* is deleterious for the individual cell, but it is beneficial for the population as a whole [12]. In general, these models include the five key properties *cell growth, formation of damage, repair of damage, cell division* and *cell death* to a varying extent. Further, a commonly implemented methodology consists of conducting simulations in order to draw conclusions about the effects of the accumulation of damage on a population level.

In this study, we take advantage of thorough mathematical analysis in combination with simulations coupled to cell growth data of single cells, in order to obtain a detailed description of the constituent parts. Moreover, we address new synergistic effects of the formation of damage, repair of damage, retention and cell size, as well as the conditions under which ageing occurs.

Here, we present a comprehensive replicative ageing model on the single-cell level. The model which includes all five key properties is validated by cell growth measurements of the increase in cell area over time for individual young and old cells. Using a mathematical methodology called non-dimensionalisation in combination with the experimental data we introduce the concept of *resilience to damage*, corresponding the difference in volume of an old and a young cell. The implemented mathematical analysis provides a deep understanding of the interplay between the fundamental constituent forces such as damage formation, repair, retention and cell size summarised in a theoretical framework describing the conditions determining if a given cell will undergo replicative ageing, starvation, immortality or clonal senescence. This framework allows us to examine ways of altering the rate of repair and formation of damage as strategies to increase the age tolerance of a single cell. Under the assumption that a minimum amount of functioning proteins is required for the cell to live, we find that asymmetric cell division enables retention of damage and that symmetrically dividing organisms cannot retain damage. Also, we show that there is a trade-off between resilience to damage of the individual cell and the capacity to retain damage. Under the same assumption, we derive the maximum degree of asymmetry that a cell can divide with as a function of the resilience to damage of the individual cell. Finally, we show the evolutionary benefit of asymmetric cell division and high damage resilience in the context of replicative ageing.

## Results

### The replicative ageing model

To investigate the interplay between the key properties underlying the replicative ageing of individual cells, we have developed a dynamic model of damage accumulation. In the model, a cell is assumed to contain two components: intact proteins and damage consisting of malfunctioning proteins. The model includes five essential properties: cell growth, formation of damage, repair of damage, cell division, and cell death (Fig 1A). Cell division and cell death are modelled as discrete events, while the dynamics of intact proteins (*P*) and damaged proteins (*D*) is continuous. The continuous part, which is described by a coupled system of *ordinary differential equations* (ODEs) (Eq (1)), is governed by cell growth, formation of damage, and repair of damage.

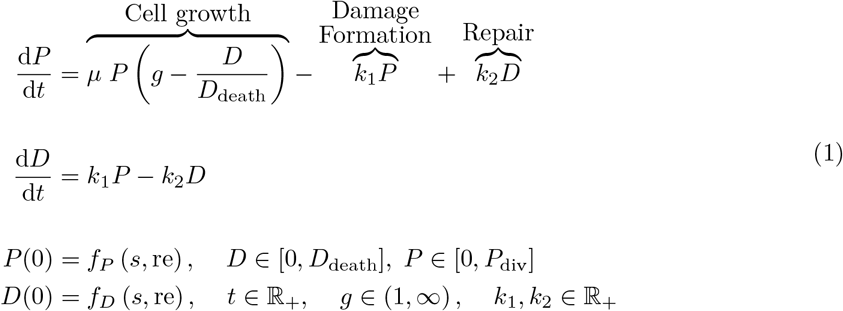

**Figure 1.**
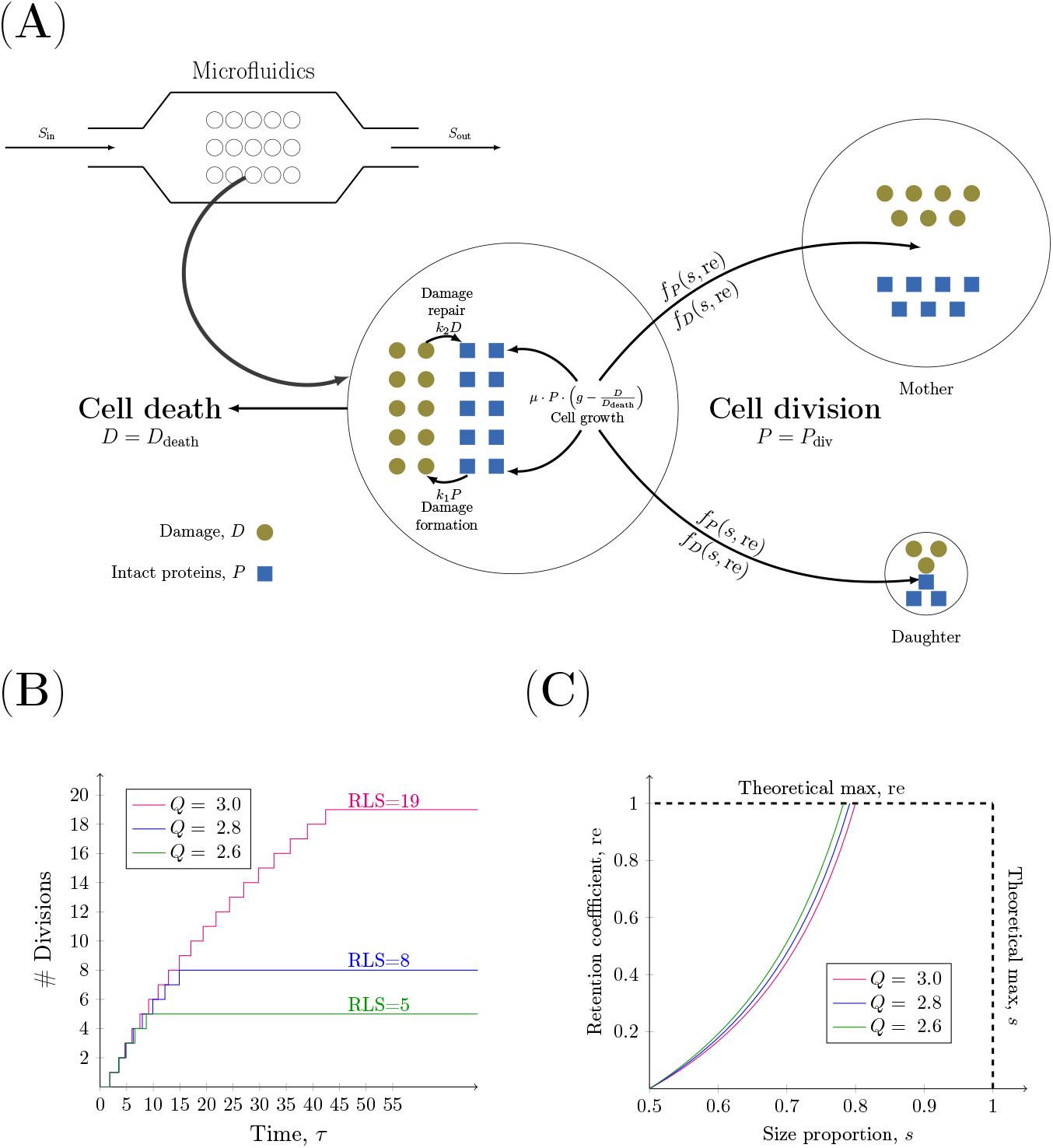
Schematic representation of the model. **(A)** The dynamics of a single cell. The production of intact proteins (blue squares) and damage (brown circles) is determined by the processes of cell growth, formation and repair of damage. When *P* = *P*_div_ cell division occurs, and the distribution of components between the mother and daughter cell is determined by the functions *f_P_* and *f_D_*. When *D* = *D*_death_ cell death occurs. Each cell is assumed to be grown in a dynamic setting such as a microfluidics device. **(B)** The effect of the damage resilience parameter *Q* = (*D*_death_/*P*_div_) on the RLS of single cells. The cell divisions are followed over time for three single cells with low [green graph : (*Q* = 2.6, RLS = 6)], medium [blue graph : (*Q* = 2.8, RLS = 9)] and high [magenta graph : (*Q* = 3.0, RLS = 20)] damage resilience. The length of the ‘’steps” of the stairs represents the generation time. The other parameters used in the simulations are *g* = 1.1, *k*_1_ = 0.5, *k*_2_ = 0.1, *s* = 0.6370, (*P*_0_, *D*_0_) = (1 − *s*, 0) and re = 0.2902. **(C)** The dependence between the degree of retention, re, and the size proportion, s. The maximum degree of retention is plotted as a function of the size proportion at the damage value *D* = 1 for three degrees of resilience to damage: low [green graph : *Q* = 2.6], medium [red graph : *Q* = 2.8] and high [magenta graph : *Q* = 3.0]

#### Cell growth

The cell growth is dictated by the availability of key nutrients, such as sugars, amino acids, and nitrogen compounds [13]. In the model, the growth of the cell is assumed to be exponential: “*μ* · *P*”. The growth rate *μ* is constant as it is assumed that an abundance of substrate is available for each cell, which would occur in a microfluidics system with continuous inflow of nutrient rich media (Fig 1A) [14, 15]. As the rate of cell growth declines with increasing amounts of damage [16, 17] the unit-less factor 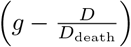 is included in the growth term. The parameter *g* is a positive number that is larger than 1 and determines the decline in growth rate. Thus, as *D* approaches the death threshold *D*_death_ the difference 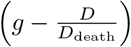 will decrease corresponding to a slower growth rate.

#### Damage formation

As a consequence of cell growth damage is formed. The various types of ageing related damage are called ageing factors or ageing determinants. They are comprised of cell compounds or cellular organelles whose functional decline over time results in a toxic effect [18, 19, 20]. In the model, we focus on damaged proteins as the ageing factor of interest. These are formed by either newly synthesised proteins that are not correctly folded or functional proteins that become unfolded. In the model, a constant proportion of the existing intact proteins *P* is converted to the reversible damage *D* with the damage formation rate *k*_1_ [*h*^−1^].

#### Damage repair

Since damaged proteins have a deleterious effect, the cell has developed several strategies to eliminate them. Damaged proteins are sorted for either repair to their proper state mediated by molecular chaperones or to degradation through the targeting of the damaged proteins to the ubiquitin-proteasome system. The system acts by moving damaged proteins into specific protein inclusions before being degraded or refolded [21, 22, 23]. In the model, we consider repair as a strategy for the cell to remove accumulated damage. A constant proportion of the existing damaged proteins *D* is converted to intact proteins *P* with the damage repair rate *k*_2_ [*h*^−1^].

#### Cell death

Cell death constitutes an important part of the process of ageing [24, 25, 26]. How the gradual deterioration over time associated with ageing ultimately leads to cell death is unknown [26]. Here, we assume that damaged proteins are deleterious for the cell and therefore cell death occurs when the amount of damaged proteins *D* reaches the death threshold *D*_death_. When this critical amount of damage is reached, the cell stops growing and is removed from the simulation. Cell division: Cell division occurs when the cell reaches a certain size. During division in *S. cerevisiae*, damaged proteins are actively retained in the mother cell [4, 19, 23]. This asymmetric segregation of damage is required to rejuvenate daughter cells and to maintain viability in populations over time. In the model, we assume that the size of a cell is proportional to the build-up of key proteins and that a cell divides when the amount of intact proteins *P* reaches the division threshold *P*_div_. Upon cell division, the intact and damaged proteins are distributed between the mother and daughter cell. This event is controlled by two parameters: the size proportion *s* and the retention value re. The size proportion 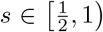 corresponds to the size of the mother cell and (1 − *s*) corresponds to the size of the daughter cell. The size proportion in the fission yeast *S.pombe* or *E.coli* is 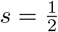 and corresponds to symmetric cell division. The bakers yeast *S.cerevisiae* divides asymmetrically with 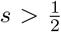 [27]. The retention value re ∈ [0, 1] corresponds to the proportion of damage that is retained in the mother cell after cell division where the value re = 1 corresponds to *all* damage being retained while no retention is given by re = 0. The distribution of intracellular components after cell division is based on the principle of *mass conservation* over generations (Eq (2)) [7]. This means that the total cellular content, that is (*P* + *D*), of the original cell before division equals the sum of the total cellular content of the mother and daughter cell after division. The conditions are also based on mass conservation with respect to intact proteins *P* and damage *D*. The initial amounts of intact proteins and damage in a cell after cell division are determined by the functions *f_P_* and *f_D_*, respectively.

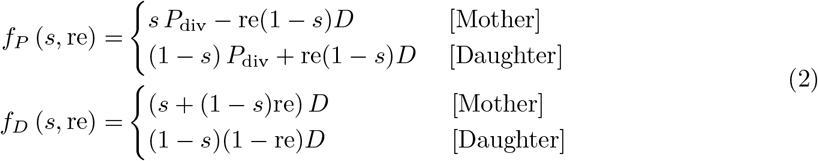

### Non-dimensionalisation introduces the property of resilience to damage

To by-pass the estimation of the parameters which can not be measured and to scale down the number of parameters, we non-dimensionalise the model. To this end, all the states and the time variable are scaled in a way such that the variable, states, and parameters of the resulting model lacks physical dimensions. This in turn simplifies the comparison between the various model components (Supplementary material S1.2). The states *P* and *D* are typically measured in molars [M] as well as their corresponding threshold values *P*_div_ and *D*_death_. Since both upper thresholds of the intact and damaged proteins are hard to estimate, we introduce the dimensionless states *P, D* ∈ [0, 1] by scaling each state with its respective threshold: *P* ← (*P*/*P*_div_) and *D* ← (*D*/*D*_death_). The introduction of these new states which are proportions of their respective thresholds results in the removal of the two threshold values *P*_div_ and *D*_death_ from the model. Similarly, the time *t* measured in hours [h] is non-dimensionalised by introducing the variable *τ* defined as *τ* = *μ* · *t*. A summary of all the dimensionless components of the model is presented in Tab 1. After the non-dimensionalisation, the continuous (Eq (1)) and discrete (Eq (2)) part of the model are given by Eq (3) and (4), respectively.

Dynamics of intact and damaged proteins:

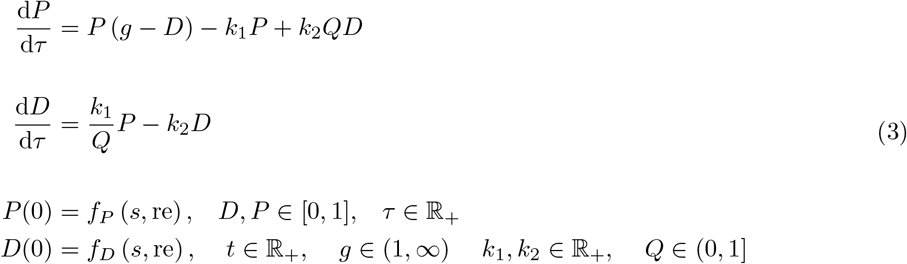

**Table 1.** The dimensionless components of the models. All the states, variables and parameters of the model are listed in the left column and their descriptions are provided in the right column.

| Component | Description |
| --- | --- |
| $P \leftarrow \frac{P}{P_{\text{div}}}$ | The proportion of intact proteins is one of the two states of the model. It lies in the interval $P \in [0, 1]$ and $P = 1$ results in cell division. |
| $D \leftarrow \frac{D}{D_{\text{death}}}$ | The proportion of damage is one of the two states of the model. It lies in the interval $D \in [0, 1]$ and $D = 1$ results in cell death. |
| $\tau = \mu \cdot t$ | The dimensionless time variable of the model is given by the time scaled by the growth rate $\mu$. |
| $g$ | A cell growth factor. It is included in the non-linear growth term “$P(g - D)$” ensuring a declining rate of cell growth with age. |
| $k_1 \leftarrow \frac{k_1}{\mu}$ | The damage formation rate as a proportion of the growth rate $\mu$ |
| $k_2 \leftarrow \frac{k_2}{\mu}$ | The damage repair rate as a proportion of the growth rate $\mu$. |
| $Q = \frac{D_{\text{death}}}{P_{\text{div}}}$ | The damage resilience parameter corresponding to the quotient between the death threshold $D_{\text{death}}$ and the division threshold $P_{\text{div}}$. A high value of $Q$ corresponds to a cell that can cope with large proportions of damage before undergoing cell death and a small value correspond to a cell that can cope only with small proportions of damage before cell death occurs. |
| $s \in [\frac{1}{2}, 1)$ | The size proportion of the daughter and mother cell with respect to the original cell. The value $s = \frac{1}{2}$ corresponds to symmetric cell division and the value $s = 0.64$ corresponds to asymmetric cell division. |
| $\text{re} \in [0, 1]$ | The retention parameter corresponding to the proportion of damage being retained in the mother cell after cell division. The value $\text{re} = 1$ corresponds to all damage being retained and the value $\text{re} = 0$ corresponds to no damage being retained. |

Cell division:

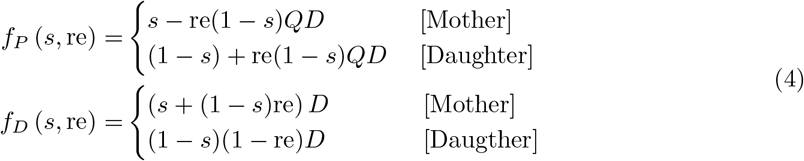

Non-dimensionalisation introduces the parameter *Q* termed the *damage resilience parameter.* It is defined as the quotient between the death and division thresholds, i.e. 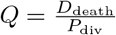, and can be interpreted as the capacity of the cell to cope with damage. A high value of *Q* corresponds to an organism that is resilient to damage with a long RLS. To test the effect of this parameter on the RLS, we have followed the number cell divisions over time for three individual cells with low, medium and high resilience to damage. As expected, an increase in the resilience to damage of a single cell yields a higher RLS (Fig 1B) and increases the maximum generation time: *τ*_max_ (*Q* = 2.6) = 6.3693, *τ*_max_ (*Q* = 2.8) = 6.9461 and *τ*_max_ (*Q* = 3.0) = 7.2382 (Supplementary Material S1.3). However, it is interesting to note that the specific generation times show the opposite trend (Supplementary Material S2.1.2). For example, the third generation time for a single cell with low resilience is longer than the third generation time for a more resilient cell. This is explained by the fact that the specific generation times in the case of low resilience are closer to the end of the life of the particular cell compared to the corresponding generation times of a more resilient cell.

### The resilience to damage can be interpreted as the difference in volume of an old cell and a young cell

It has been reported that mother cells are clearly distinguished from their daughters as their size increases steadily with successive divisions where an old cell corresponds to a large cell in terms of volume and mass [28, 3, 29, 30, 31, 32]. These experimental studies suggest that the average volume of old cells at the end of their life is approximately 3.5 times larger than that of virgin daughter cells born with no damage at the point of cell division, which also is observed in our experiments of dividing young and old cells (Supplementary material S1.1).

As each cell consists of intact proteins *P* and damage *D* (Fig 1A), it is reasonable to assume that the total protein content is proportional to the cell size, which can be approximated as an area (i.e. cell area ∝ *P* + *D*). Typically, this is given by measuring the cell area obtained by time-lapse microscopy imaging [31]. In the context of the dimensionless model, the corresponding output can be written as *ŷ*(*θ, τ*) = *P* (*θ, τ*) + *QD* (*θ, τ*), where *θ* = (*g*, *k*_1_, *k*_2_)^*T*^ is the parameter vector consisting of the involved rate parameters and *τ* is the dimensionless time. This indicates that the damage resilience quotient *Q* corresponds to the increase in size of an old cell compared to a young cell. Assuming that a daughter cell born with no damage has accumulated almost no damage at the point of budding (i.e. *D* ≈ 0), its dimensionless area is *y* ≈ 1 as cell division occurs when *P* =1. An old mother cell undergoes cell death when *D* =1 and hence the volume of this cell at the point of cell death is *y* = 1 + *Q*. Accordingly, the damage resilience quotient should be *Q* ≈ 2.5. It is of interest to note that damage resilience is embedded in the proposed modelling framework for describing replicative ageing and that it is independent of the specific dynamics assumed in the model. As the property is introduced by the non-dimensionalisation procedure, the formulation of the ODE’s is independent of the damage resilience parameter. In other words, using the proposed non-dimensionalisation the property of resilience to damage will be introduced independent of the assumptions made on the forces cell growth, formation and repair of damage.

### Model validation

To evaluate the performance of the model and its reliability it is fitted to experimentally obtained data of dividing young and old cells (Fig 2 and Supplementary material S1.1). Further, we compare this model with two selected models that explicitly focus on the accumulation of damage in budding yeast [10, 7] (Supplementary material S1.2 and S1.2.2 and S1.2.3).

**Figure 2.**
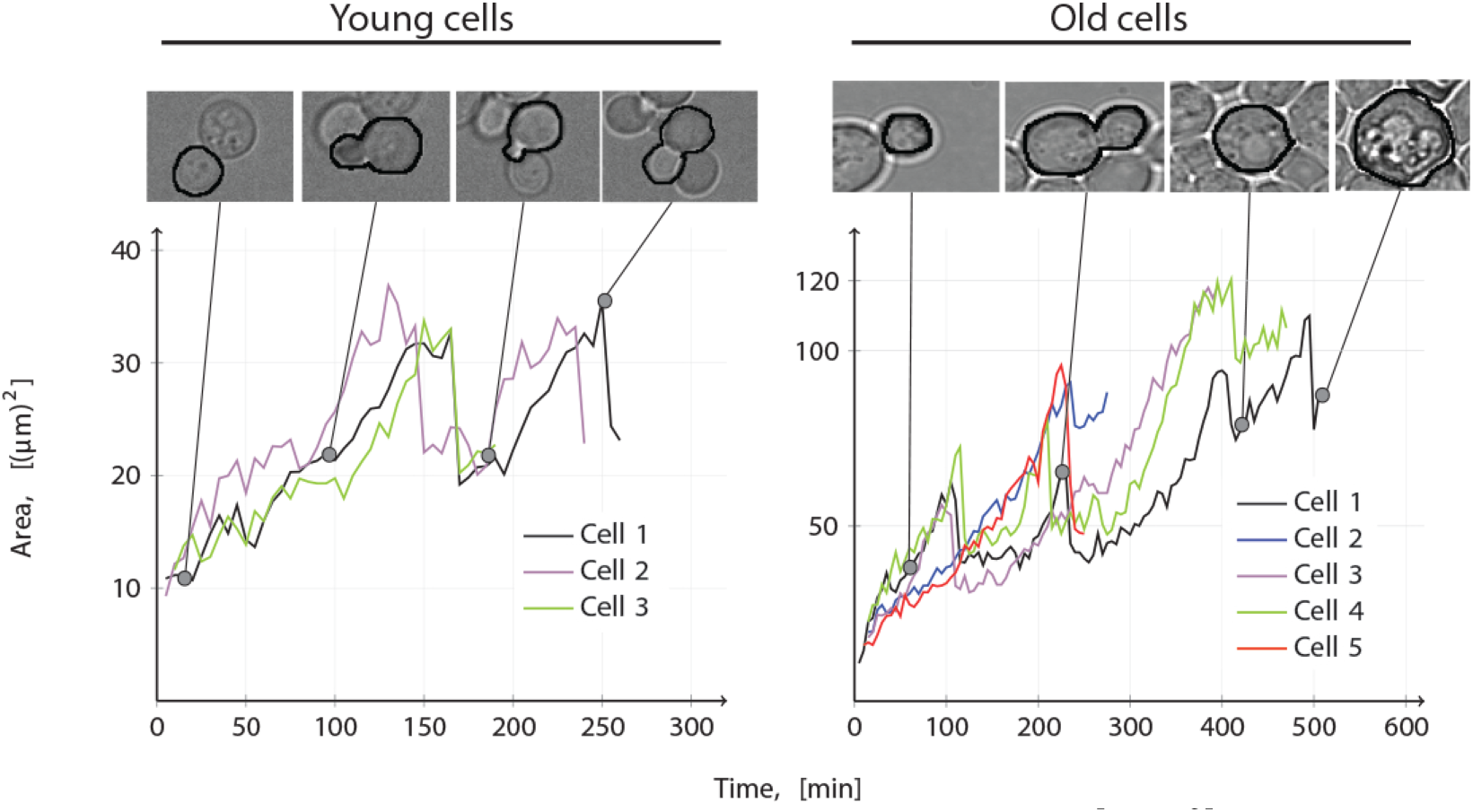
Time series data of cell area over time. The cell area [(*μ*m)^2^] of individual wild type yeast cells is plotted over time [min]. The left hand figure shows the increase in cell area for three “damage-free” daughter cells and the right hand figure shows the corresponding increase in cell area for five old mother cells.

As a representation of the total protein content on a single-cell level, we measure the cell area obtained from bright-field microscopy images (Material and Methods). These images are taken of both young and old yeast cells (Supplementary material S1.1). To enable a continuous availability of nutrients required for cell growth and division the cells are grown in a microfluidics device under a continuous inflow (and outflow) of media, ensuring that the cells are exposed to optimal growth conditions regarding nutrients. With our setup we can observe the cell area, as a measurement of the total protein content, and follow the growth and division of a single cell over time. Therefore, we use this setup to obtain time series measurements of cell area of young and old cells for at least 1 division per cell (Fig 2).

The model validation shows that in terms of both the least square (LS) value of the fit and the *Akaike information criterion* (AIC), which accounts for the model complexity, the presented model ((LS, AIC) = (0.20, −288)) outperforms both the model by Erjavec et al.[7] ((LS, AIC) = (0.46, −210)) and the model proposed by Clegg et al.[10] ((LS, AIC) = (0.43, −219)) (Supplementary material S1.4). Both these numbers should be as low as possible, where the latter criterion suggests that the model with the least parameters in combination with the best fit should be selected [33].

Further, as the replicative ageing is characterised by an increase in cell size [28, 3, 29, 30, 31, 32], for the models to describe replicative ageing they should satisfy the criteria that RLS ∝ *Q* independent of the rate parameters that are selected. Thus, by picking parameters giving rise to a finite RLS, an increase in Q while keeping the remaining parameters fixed should increase the life span, while a decrease in Q should decrease the life span. The simulations show that our model together with the model by Erjavec et al. [7] satisfy this criteria which the model by Clegg et al.[10] does not, and hence our theoretical description captures this important aspect of replicative ageing in yeast (Supplementary material S1.4).

This finding motivates further investigation of the properties of the presented model as it can describe ageing in yeast correctly. The subsequent results are the outcome of the mathematical analysis of the discrete (Eq (4)) and continuous (Eq (3)) parts of the model.

### Asymmetric division allows for retention of damage which comes at the price of a lower resilience to damage

Using the presented theoretical framework, it is of interest to see how retention of damage by the mother cell is influenced by the other factors of the model. More specifically we address the following questions: (1) how much damage can a mother cell retain, (2) how does the capacity to retain change throughout the life time, (3) what factors limit the amount of damage a mother cell can retain at cell division and (4) how does the degree of asymmetry in the cell division affect the capacity to retain damage.

Assuming that the minimal amount of intact proteins that a cell is required to have after cell division is *P*_0,min_ = (1 − *s*)P_div_ or *P*_0,min_ = (1 − *s*) in the dimensionless case, we derive a mathematical constraint (Supplementary material S2.2.1) addressing the above considerations (Eq (5)). This minimal amount of intact proteins is required for the cell to grow, perform vital cellular activities and subsequently divide. The lower limit is based on the fact that the smallest unit of life for unicellular systems according to the assumptions of the model is a damage free daughter cell with the initial conditions (*P*_0_, *D*_0_) = (1 − *s*, 0), and to grow it needs a proportion of at least *P*_0,min_ = (1 − *s*) intact proteins initially. Moreover, assuming that a mother cell can maximally retain damage so that it has at least *P*_0,min_ intact proteins after cell division, it is possible to derive constrains on how much damage a cell can retain at the point of cell division (Eq (5)).

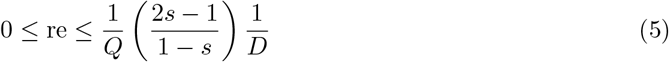

Three main conclusions can be drawn from the upper constraint on the retention of damage. Firstly, the capacity to retain damage is inversely proportional to the amount of damage that the cell contains, i.e. re ∝ 1/*D*, implying that the capacity to retain damage decreases as the amount of damage increases.

Secondly, the capacity to retain damage is inversely proportional to the degree of resilience, i.e. re ∝ 1/*Q*. This result suggests that investing resources in the capacity to retain damage comes at the cost of a lesser degree of resilience to damage for the individual cell.

Thirdly, retention is a byproduct of asymmetric division. In the case of symmetric division, i.e. *s* = 1/2, the upper limit vanishes (Eq (5)) and thus there is no damage retention, re = 0. Furthermore, it holds that the maximum degree of retention is proportional to the degree of asymmetry, i.e. re ∝ *s*. This theoretical description illustrates the dependence between the retention coefficient on the one hand and the size proportion and damage resilience on the other (Fig 1C). Moreover, the maximal proportion that a cell can retain is re = 1 which allows us to derive a condition for the *maximum degree of asymmetry* at which a cell can divide (Eq (6)).

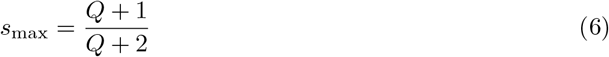

Given the value *Q* = 2.5 of damage resilience, which has been observed by us and others [29, 31, 32], the expected maximal degree of asymmetry is *s*_max_ = 0.8.

Mathematical analysis of the discrete part of the model (Eq (4)) resulted in two equations (Eq (5) and (6)) connecting all the important parameters linked to the cell division, namely s, re and *Q*. In a similar manner, it is of interest to understand how the rate parameters for the formation of damage *k*_1_ and repair of damage *k*_2_ controls the dynamics of the continuous part of the model (Eq (3)).

### The conditions allowing for replicative ageing

In order to pick biologically relevant parameter pairs (*k*_1_, *k*_2_), a condition based on nutrient availability is imposed on the model. Given enough food in the system, the cells should grow and as a consequence of cell growth damage is accumulated within the cell [34]. This implies that both states *P* and *D* should be increasing functions of time and this is ensured by using linear stability analysis of the steady states (Supplementary material S2.2). Using the above approach, we define the conditions that allow for replicative ageing, resulting in a theoretical framework classifying all possible types of dynamics for any cell into four categories: *starvation*, *immortality*, *ageing*, and *clonal senescence* (Fig 3A). These correspond to four different regions of the parameter space, which are defined by *starvation, immortality* and *clonal senescence* constraints.

**Figure 3.**
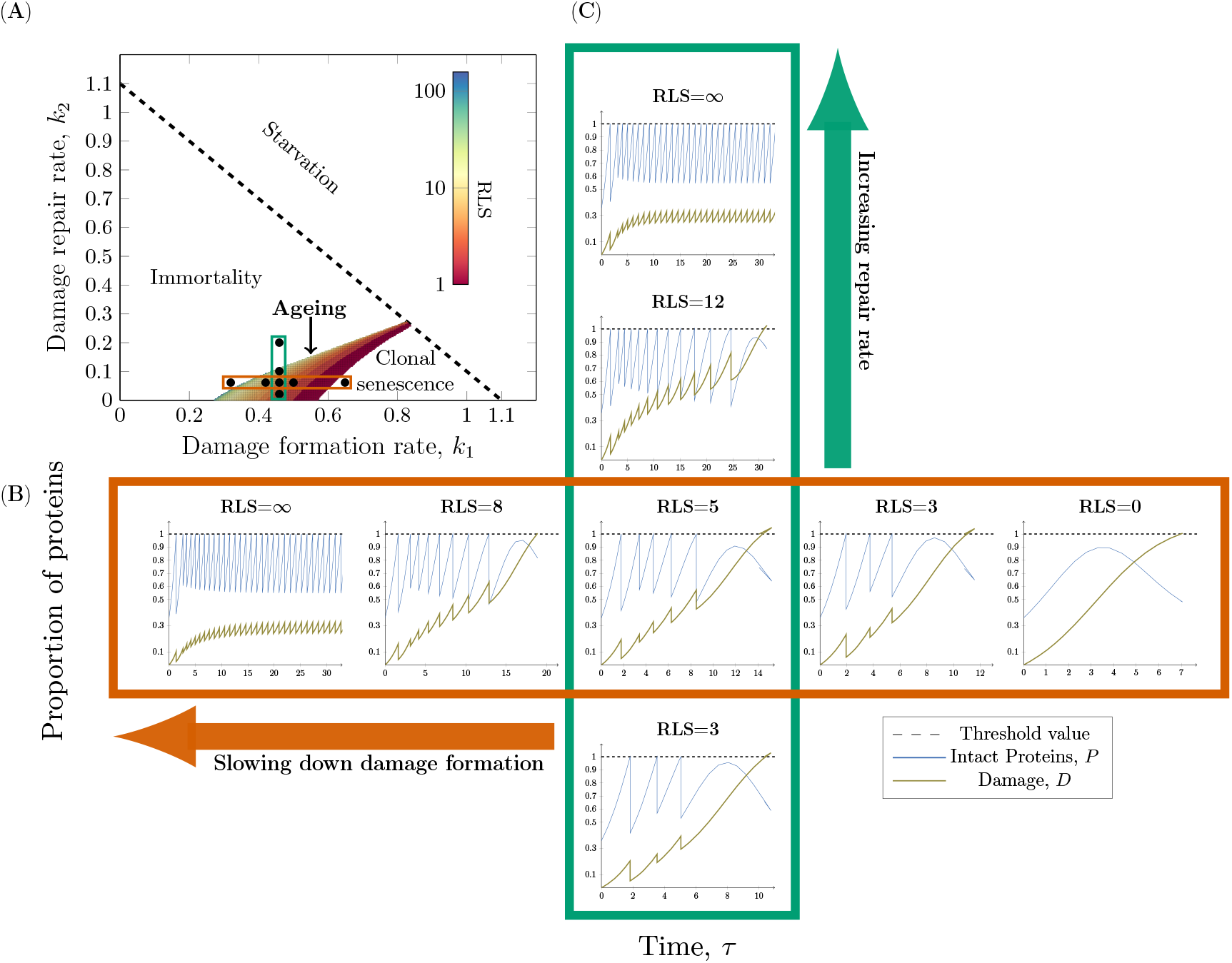
The interplay between damage formation and damage repair. **(A)** The constraints on the rate of damage formation *k*_1_ and repair *k*_2_ define four different regions: *Starvation*, *Immortality*, *Clonal senescence* and *Ageing*. Cells with parameters above the dashed line will consume substrate too quickly and thereby undergo Starvation. Below the dashed line the parameters correspond to three types of cells characterised by Immortality (infinite RLS), Ageing (finite RLS) and Clonal Senescence (no RLS). Within the ageing region of the parameter space the replicative life span (RLS) corresponding to a set of parameters is presented in the colour bar. (**B** & **C**) The formation of intact proteins *P* and damage *D* is simulated over time *τ* until cell death occurs. The threshold value determines when cell division (*P* = 1) or cell death (*D* = 1) occurs. **(B)** *Decreasing the formation of damage increases the RLS* (*k*_1_ ∈ {0.32, 0.42, 0.46, 0.50, 0.65} and fixed repair rate *k*_2_ = 0.06). **(C)** *Increasing the repair rate increases the RLS* (*k*_2_ ∈ {0.02, 0.06, 0.1, 0.2} and fixed damage formation rate *k*_1_ = 0.46). The other parameters used in the simulations are *g* = 1.1, *Q* = 2.6, *s* = 0.64, (*P*_0_, *D*_0_) = (1 − *s*, 0) and re = 0.299.

The *starvation* constraint predicts the minimum amount of substrate required for growing and therefore ageing (Eq (7)). Cells with parameters within the starvation region will not be able to form sufficient amounts of proteins leading to a collapse of the cellular machinery. The starvation constraint connects the uptake of nutrients to the damage accumulation process by acting as an upper bound on the sum of the damage formation and repair rates. The critical amount of substrate necessary for an organism to undergo ageing is 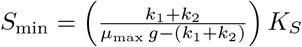 in the case of Monod growth (that is when 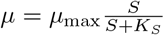) and if this constraint is not satisfied the cell will undergo starvation.

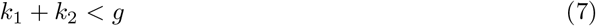

Cells that do not satisfy the *immortality constraint* have an infinite RLS. It constitutes a lower bound on the damage formation rate or equivalently an upper bound on the damage repair rate.

Cells that do not satisfy the *clonal senescence constraint* have a RLS of zero divisions implying that they do not divide before undergoing cell death. It acts as an upper bound on the damage formation rate or a lower bound on the repair rate

The *ageing* region represents every cell within the population that has a *finite* replicative life span implying that it should divide at least once and die after a finite number of cell divisions [3]. The RLS of an individual cell in the ageing region is inversely proportional to its rate of damage formation and proportional to its rate of repair which, enables the construction of strategies for prolonging the RLS.

Next, we investigate the effect of the model parameters on the RLS of individual cells. As the formation and repair of damage are fundamental to the accumulation of damage, a profound understanding of these forces is required in order to fully grasp the mechanisms behind replicative ageing. Within the ageing region, the RLS can be increased by two obvious strategies: by decreasing the damage formation rate *k*_1_ (Fig 3B) and by increasing the damage repair rate *k*_2_ (Fig 3C). Here, we present the dynamics of the intact and damaged proteins for eight different damage-free daughters to illustrate the effect of altering the rate parameters according to the two proposed life prolonging strategies (Fig 3).

As expected, the results of the simulations show that a decrease in the formation of damage affects the RLS of a single cell (Fig 3B). Similarly, an increase in the repair efficiency increases the RLS as well (Fig 3C). Next, we set out to explore synergistic effects between retetnion of damage, formation of damage and damage repiar on increas of RLS, by systematicaly analysing a large set of model parameters.

### The role of retention, damage formation and repair in replicative ageing

The strategies for prolonging the RLS are generalised in two steps. Firstly, the effect of retention on the two strategies is added to the analysis and secondly, the gains in RLS from the two strategies are compared for numerous cells with different capacity to retain damage. In our model, the distribution of damage depends on both the cell size and the damage retention parameter. Therefore, we compare the cases of no (re = 0, the mother effectively retains 64% of the existing damage) and high (re = 0.299, the mother effectively retains 74.8% of the existing damage) retention. The results show that the ageing area of the parameter space increases with retention at the expense of the immortality region (Fig 4A).

**Figure 4.**
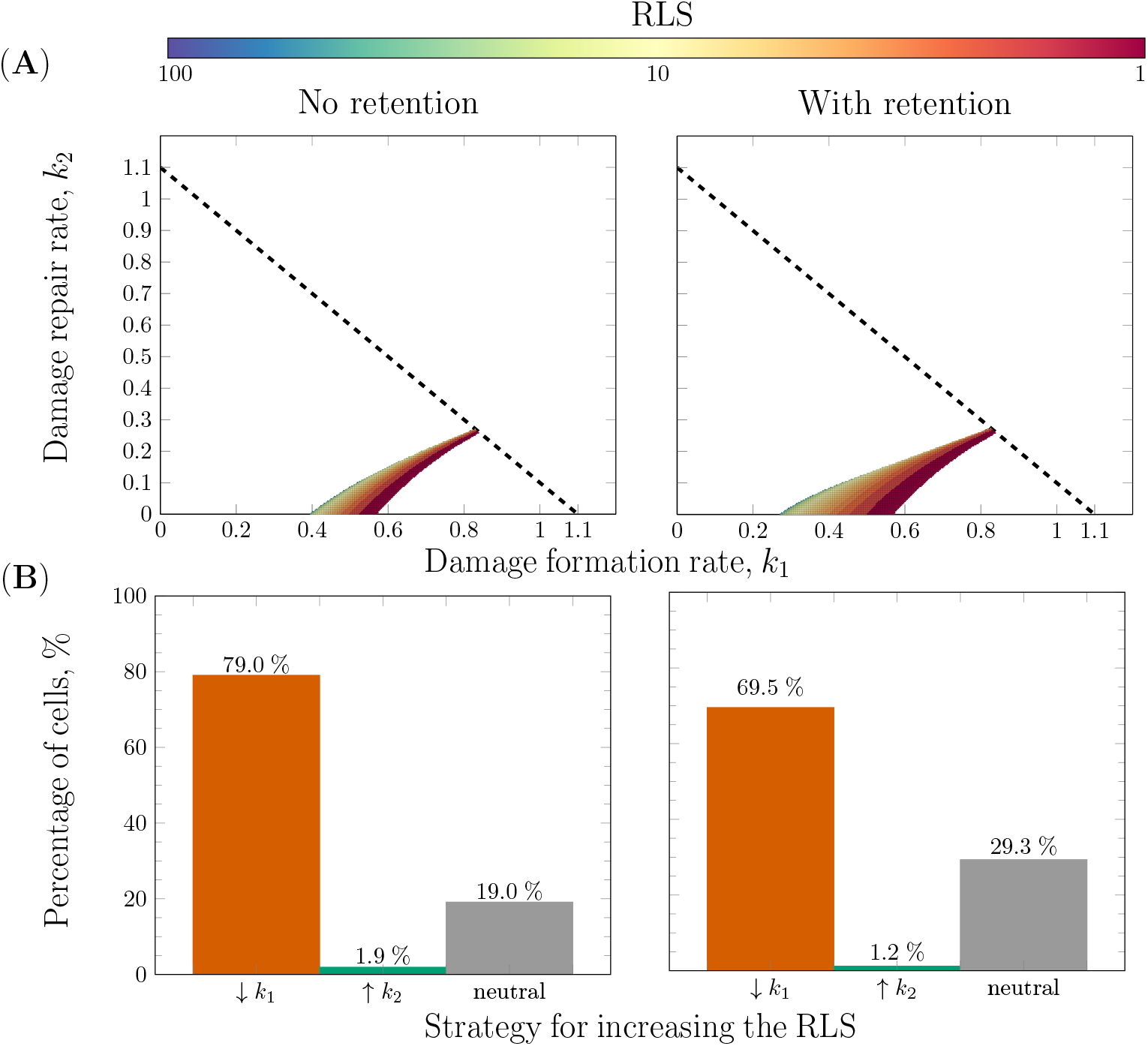
The effects of retention and repair on replicative ageing.. **(A)** The effect of retention on the ageing area with two retention profiles: no retention re = 0 on the left and with retention re = 0.299. The area of the ageing region increases to the left at the expense of the immortality region proportionally to the degree of retention of damage. **(B)** The efficiency of the RLS prolonging strategies as a function of retention. The increases in RLS of single cells by increasing the repair rate and decreasing the formation of damage are compared for the same two retention profiles as in (A). The cells are divided into three categories: Orange (‘’↓ *k*_1_”): *decrease in formation of damage* where ΔRLS_*k*_1__ > ΔRLS_*k*_2__, Green (‘’↑ *k*_2_”): *increase in rate of repair* where ΔRLS_*k*_1__ < ΔRLS_*k*_2__ and Grey (‘’neutral”): *both strategies* are equally good where ΔRLS_*k*_2__ = ΔRLS_*k*_2__. The other parameter used in the simulations are *g* = 1.1, *Q* = 2.6, *s* = 0.64 and (*P*_0_, *D*_0_) = (1 − *s*, 0).

Next, we compare the two strategies for increasing the RLS of single cells with and without retention (Fig 4B). The life spans of numerous damage-free daughter cells with equidistant parameters are simulated. More precisely, according to the starvation constraint (Eq (7)) the parameters for the rate of formation and repair of damage lies in the following interval: *k*_1_, *k*_2_ ∈ [0, *g*]. Moreover, if the interval [0, *g*] is partitioned into *N* sub intervals (the integer *N* ∈ ℤ^+^ defines the mesh size) with equal lengths, then the length of each interval is 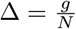. Using a specific mesh size *N* × *N*, it is possible to loop over the (*k*_1_, *k*_2_)-parameter space and generate the ageing landscape with different degrees of accuracy determined by the mesh size. At each point in this grid, the notation Δ_*k*_1__ corresponds to decrease in damage formation with 5% and a corresponding increase of 5% in repair is denoted Δ_*k*_2__. Thereafter the gain in RLS by decreasing the formation of damage given by ΔRLS_*k*_1__ = RLS (*k*_1_ − Δ_*k*_1__, *k*_2_) − RLS (*k*_1_, *k*_2_) is compared to the gain in increasing the repair given by ΔRLS_*k*_2__ = RLS (*k*_1_, *k*_2_ + Δ_*k*_2__) − RLS (*k*_1_, *k*_2_). Provided these values, all parameters for the two retention profiles are divided into three categories: *increasing the rate of repair* ‘’↑ *k*_2_” where ΔRLS_*k*_1__ < ΔRLS_*k*_2__, *decreasing the formation of damage* “↓ *k*_1_” where ΔRLS_*k*_1__ > ΔRLS_*k*_2__ and the *neutral strategy* ‘’neutral” where ΔRLS_*k*_1__ = ΔRLS_*k*_2__ (Fig 4B).

We found that for the majority of cells, ca 79% without retention and 70% with retention, decreasing the formation of damage is a more efficient RLS-prolonging strategy (Fig 4B). Moreover, for strains with a functioning retention mechanism the number of cells for which both strategies are equally efficient is higher compared to the corresponding number for strains without retention, namely ca 29% in the former case compared to 19% in the latter.

### Asymmetric cell division and high resilience promote higher replicative lifespan

We evaluate the significance of asymmetric division and resilience to damage by comparing symmetric (*s* = 0.50) and asymmetric (*s* = 0.64) cell division (with and without retention) under low (*Q* = 2.5) and high (*Q* = 3.0) resilience to damage (Fig 5). An increase in resilience to damage shifts the ageing region to the right implying that resilient organisms undergo ageing at high rates of formation of damage. The effect of an increased resilience is that the RLS of a cell is prolonged since the immortality region moves closer to the corresponding point in the parameter space as the value of *Q* increases. This also confirms the previous simulations (Fig 1B) and (Fig ??).

**Figure 5.**
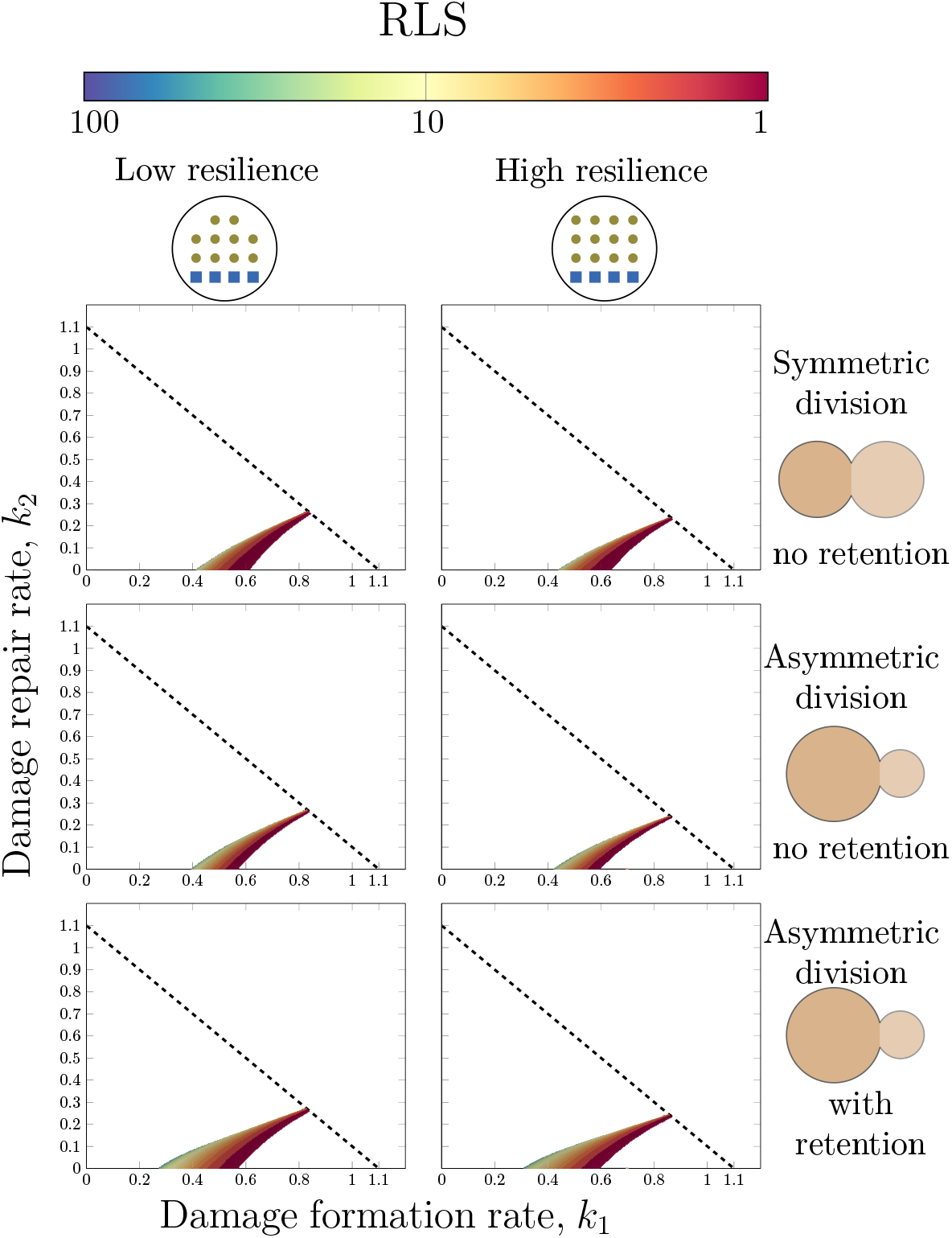
The benefit of asymmetric cell division and a high resilience to damage in the context of ageing. Two levels of resilience to damage (*Q* = 2.6 left column & *Q* = 3.0 right column) and three different size proportions in the cell division (top row: (*s*, re) = (0.5, 0), middle row: (*s*, re) = (0.64, 0) & bottom row: (*s*, re) = (0.64, 0.2593) in the left sub figure and (*s*, re) = (0.64, 0.3111) in the right sub figure). The other parameters used for are *g* = 1.1 and (*P*_0_, *D*_0_) = (1 − *s*, 0).

Similarly, a higher degree of asymmetry in the cell division yields a longer life span. The RLS is higher in the asymmetric (middle row of Fig 5) compared to the symmetric (top row of Fig 5) case giving weight to the proposed link between asymmetric cell division and longevity in the context of replicative ageing. This is supported by the comparison between stem cells for which the asymmetric cell division results in a rejuvenated daughter cell and symmetrically dividing neurons where the amount of damage is greater due to the incapacity to remove damage through cell division [35]. As retention is added, the ageing area increases (middle vs bottom row of Fig 5) and also the RLS is higher in a larger part of the ageing region of the parameter space for the asymmetrically dividing organisms with retention than the counterpart without. As we showed previously, asymmetric division enables retention of damage and this result indicates that retention and ageing are closely interlinked.

## Discussion

To achieve a comprehensive understanding underlying the phenomena of ageing in unicellular systems, the interplay between the formation, repair, resilience and distribution of damage upon cell division is vital. In this study, we developed and validated a mathematical model of replicative ageing accounting for all key properties: *cell growth*, *formation of damage*, *repair of damage*, *cell division* and *cell death* (Fig 3A), defining the conditions leading to starvation, immortality, clonal senescence and ageing. This detailed theoretical analysis and description of the model properties resulted in five main findings: (I) The capacity to retain damage decreases as the amount of damage increases (Eq (5)). (II) There is a trade off between damage resilience and damage retention (Eq (5)). (III) There is a maximal degree of asymmetry at which a cell can divide governed by the introduced property resilience to damage (Eq (6)). (IV) Retention of damage is a byproduct of asymmetric division and is closely connected to replicative ageing (Fig 4A). (V) Asymmetric cell division and damage resilience are tightly coupled to replicative ageing (Fig 5).

The presented model is validated using time-lapse microscopy data of cell area over time for both young and old cells (Fig 2). It is novel in three respects: it links all major biological features of accumulation of damage, it focuses on replicative ageing and its methodology combines mathematical analysis with simulations. As a consequence of the non-dimensionalisation procedure, the intrinsic property of damage resilience *Q* emerges which is essential to the replicative ageing of a single cell where an increase in resilience greatly prolongs the life span (Fig 1C & 5). Our model is in agreement with previous results showing the critical role of asymmetric segregation of damage through retention in the context of ageing [11, 12] and that cells forming damage with a rate under a certain threshold result in immortal lineages [11]. In addition, our framework connects all key factors, having profound implications on replicative ageing, and it enables the study of the relationships between properties of ageing as opposed to studying them individually. For example, we show that the starvation constraint (Eq (7)) connects the uptake of food to the damage accumulation process.

The results further indicate that the retention mechanism is tightly coupled to replicative ageing (Fig 4A) and it allows the cell to reach a higher RLS in a more robust fashion (Fig 4B). The cells with a functioning retention mechanism undergo ageing at lower rates of formation of damage. Besides, a high degree of retention increases the set of parameters yielding a high RLS implying that retention is also fundamental for longevity. This reaffirms the importance of the budding event with respect to the ageing process in *S.cerevisiae*. We also observe that in order to increase the RLS the cells should invest resources in decreasing the damage formation rate, regardless of the capacity to retain damage (Fig 4B). However, for cells with a functioning retention machinery an increase in the efficiency to repair damage prolongs the RLS more compared to a similar increase for cells without retention.

Moreover, we propose a precise relationship between damage resilience, damage retention and cell size (Eq (5)). This has been suggested in previous experimental studies [36], however how these three forces depend on each other remain elusive. For an average yeast cell with a damage resilience of *Q* = 2.6, the maximal degree of asymmetry corresponds to a size proportion of *s* = 0.8. Our data suggest that the budding yeast divides with *s* = 0.64 (Supplementary material S2.1.1), which allows for an optimal trade-off between damage resilience and damage retention, where a high resilience to damage corresponds to a cell that can obtain a high age and retention corresponds to a sacrifice of fitness for the individual cell for the sake of its offspring. Thus, (Eq (5)) describes the balance between investing in altruism versus selfishness in the context of ageing. Furthermore, the capacity to retain damage deteriorates with high age which can be motivated by the fact that the functions of numerous cellular processes decline during the ageing process. Considering that the maximal degree of asymmetry is proportional to damage resilience (i.e. *s*_max_ ∝ *Q*), it implies that the higher the resilience or equivalently the larger the cell volume the more asymmetric the cell division can be (Eq (5).

Despite the fact that several parameters of the model are unidentifiable, the good fit to the experimental data and its ability to capture the replicative ageing process indicate the correctness of the model structure. Moving forward, the quantifiability of the model can be increased by considering each of the key processes as modules. More precisely, by defining the modules cell growth, repair, formation, cell division and cell death it is possible to expand the level of detail in each module by studying specific intracellular pathways connected to these processes. Using the current model as interfaces between these modules it is possible to construct a quantitative model of replicative ageing in unicellular organisms like *S.cerevisiae*.

The proposed theoretical framework suggests that asymmetric division is beneficial in the context of replicative ageing (Fig 5). When comparing the effect of symmetric versus asymmetric cell division, our results indicate that the latter mode of division yields a higher RLS compared to the former. In order to ensure the viability of cell lines, the asymmetric segregation of damage at the point of cell division is crucial. This segregation is ensured by the retention mechanism but it has also been speculated that it is enhanced by asymmetric division. When comparing asymmetrically dividing stem cells with symmetrically dividing neurons the former class of cells is able to remove damage through cell division to a larger extent while the latter has a higher instance of damage inclusions [35]. We also show that an organism dividing purely symmetrically cannot have an active damage retention mechanism (Eq (5)). In addition, it has been shown that the short-lived *sir2*Δ mutant of yeast divides more symmetrically and fails to segregate damage [36].

Further, we show the link between cell growth and formation of damage. Under favourable conditions in terms of available substrate, it has been speculated that *E.coli* does not undergo replicative ageing [37, 38, 8] which entails that its parameters are located in the immortality region of the parameter space (Fig 3A). Such parameters correspond to low rates of formation of damage which in turn corresponds to a high growth rate as *k*_1_ ∝ (1/*μ*) (Tab 1). Furthermore, the genome size of *E.coli* is substantially smaller compared to that of *S.cerevisiae*, namely ~4.64 Mbp [39] versus ~12.07 Mbp [40]. Due to the small genome size, it is not required to spend large resources on the maintenance of the cell. From this follows that it is reasonable to assume that *E.coli* has a low rate of formation of damage. In accordance with the “Disposable Soma”-theory [41], this allows it to prioritise cell growth over maintenance, and thus a high rate of cell growth should be indicative of a low rate of damage formation which is what our model suggests. Besides, it has been shown that *E.coli* has optimised its metabolic flux to achieve a maximal growth rate [42]. This suggests that *E. coli* strongly favours growth, and therefore spends less energy on maintenance which results in it not undergoing ageing when growth conditions are optimal. Furthermore, Wasko et al. proposed that growth rate, rather than replicative lifespan, is a more appropriate measure of fecundity [43]. They argue that there is a minimal benefit of increasing the lifespan by one generation on the population level in contrast to the significant gain brought by a small increase in growth rate. These observations establish the close link between cell growth and formation of damage. This implies that rapidly growing organisms with small genomes do not undergo ageing under optimal conditions. On the other hand, slower growing organisms with larger genomes are forced to maintain their cell and therefore undergo replicative ageing as a by-product of growth.

This balance between growth and maintenance of the cell is captured by the proposed model. Thus our work raises the question if ageing is a consequence of the necessity of cell maintenance for certain organisms in specific environmental conditions.

## Materials and Methods

### Time-lapse microscopy and cell tracking

The yeast cells (BY4741, MATa his3Δ1 leu2Δ0 met15Δ0 ura3Δ0) were inoculated in 5 ml of complete synthetic medium media containing 1.7 g/l yeast nitrogen base, 5 g/l ammonium sulfate, 670 mg/l complete supplement mix supplemented and 4 % glucose from freshly streaked (2-4 days old) plates and incubated in a shaker at 30°C for at least 8 hours on the day before the experiment. These cell were injected with a syringe in a two-channel Y-formed microfluidics poly-dimethylsiloxane (PDMS) system and allowed to sediment in the main channel. Fresh CSM media was supplied to the cells through the other channel. The experimental setup is further described in Welkenhuysen et al. 2018[44]. Imaging was performed on a Leica DMi8 inverted fluorescence microscope (Leica microsystems). The microscope was equipped with a HCX PL APO 40x/1.30 oil objective (Leica microsystems), Lumencor SOLA SE (Lumencor) led light and Leica DFC9000 GT sCMOS camera (Leica microsystems). Cell growth was recorded at 1 frame in bright-field at 20 ms exposure every 5 minutes. Only daughters who have been budded of from the mother cell in the PDMS system were observed for their first divisions. Analysis of cell area was performed with the ImageJ distribution FIJI [45].

### Numerical implementation

All the numerical results, except for the structural identifiability analysis, were generated using Matlab [46]. The system of ODEs for the various models have been solved using the built-in solver `ode45` which uses an adaptive “Runge-Kutta-Fehlberg”-method of order 4 and 5 [47]. The model validation have and the numerical identifiability analysis have been conducted using the optimiser `lsqnonlin`. The structural identifiability analysis of the three competing models have been obtained using the Mathematica software developed by Karlsson et al.[48]. A detailed description of the numerical implementations of the algorithms used for generating the results can be found in the Appendix (See Supplementary material S3).

## Supporting information

**S1 Appendix. Supplementary material**. The appendix contains the data, the parameter values used in the study, a detailed mathematical analysis and a description of the numerical implementations.

## Author contribution

JB developed the mathematical model, performed simulations and mathematical analysis. NW planned and performed experimental part of the work (microscopy and microfluidics experiments). MC conceived the research. JB and MC designed the research. JB, NW and MC wrote the paper.

## Acknowledgments

This work was supported by Swedish Agency for Strategic Research (grant nr. IB13-0022) and Hasselblad foundation. We would like to thank all past and present members of the CvijovicLab for valuable input and careful reading of the manuscript.

