## Supplementary Material for "Repair, resilience and asymmetric segregation of damage in the context of replicative ageing: it is a balancing act"

| | |
|---|---|
| *Scripts and data-files*: | `https://github.com/cvijoviclab/Ageing.git` |
| *Raw data (microscopy images)*: | `https://figshare.com/s/241d6664766ac8b5b2b9` |

Johannes Borgqvist[1], Niek Welkenhuysen[1], Marija Cvijovic[1*]

[1] Department of Mathematical Sciences, Chalmers University of Technology and the University of Gothenburg, Gothenburg, Sweden.

*Corresponding Author: Marija Cvijovic, Chalmers University of Technology and the University of Gothenburg, Mathematical Sciences, Chalmers tvärgata 3, SE-412 96 Gothenburg, Sweden Phone: +46 31 772 53 21, e-mail: ``

### S1 Data, Parameter Estimation & Model validation

#### S1.1 Data of cell growth for individual cells of *S.cerevisiae*

For the purpose of validation, the model is fitted to experimental data of cell growth (Fig S1). Using microscropy, the increases in cell area $A$ $\left[(\mu\mathrm{m})^2\right]$ over time $t$ [min] have been measured for numerous both young and old wildtype cells of *S.cerevisiae*. As the model is qualitative, it describes the dynamics of intact and damaged proteins regarded as bulk quantities. Thereby, the model describes overall features of the cell as oppose to physical quantities, and in order to conduct the validation both the data and the model must be rendered dimensionless. For both the modelling output and the experimental data, the "area" and the time must be rendered dimensionless.

To exemplify the scaling procedure, the left hand figure displays the raw data of a growing daughter cell (Fig S1). The studied daughter is young in the sense that it is born without damage, and in the raw data it grows until cell division occurs which corresponds to the "jumps" in the data where the mother cell looses mass due to the formation of a daugther cell before continuing growing. Hence, using the cell mass before and after the jump, the size proportion parameter $s$ can be estimated from the data by taking the quotient of the upper and lower data point, i.e. $s_1 = \dfrac{y_1}{y_2}$ where the subscript 1 corresponds to the first cell division to use the notation introduced in Fig S1. By taking the average of multiple "jumps", i.e. multiple cell divisions, for multiple cells it is possible to estimate the mean size proportion $s$ which is the one that has been implemented in the model validation procedure. Remarkably enough, the jump size is rather constant which thus validates the assumption that the mother cell obtains a proportion $s$ of the cell mass before division while the daughter obtains the corresponding proportion of $(1-s)$.

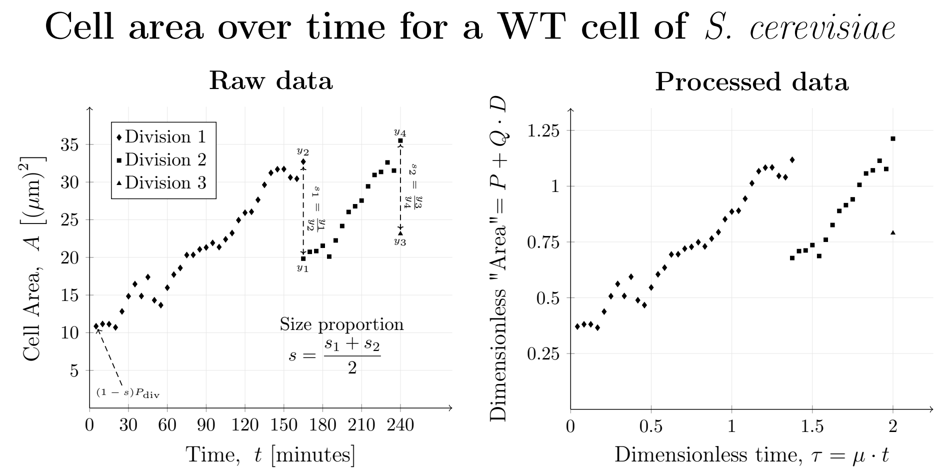

**Figure S1. Non-dimensionalisation of data of the increase in cell area over time for a single daughter cell of *S.cerevisiae*.** The increase in cell area over time for a wild type cell of *S.cerevisiae*. The left hand figure corresponds to the raw data and the right hand figure corresponds to the processed dimensionless data. The processed data is obtained by scaling the cell area on the $y$-axis by the value $(P_{\mathrm{div}})^{-1}$ which is estimated from the data and the time on the $x$-axis is rendered dimensionless by scaling it by $\mu = 0.5\ \mathrm{h}^{-1}$. The value of $P_{\mathrm{div}}$ is calculated by assuming that the first data point equals the value $(1-s)\cdot P_{\mathrm{div}}$ where $s$ is the size proportion. The size proportion $s$ is calculated by the two "jumps" associated with each cell division.

Using the size proportion $s$, it is possible to estimate the value $P_{\mathrm{div}}$ used to scale the area. The estimation of the parameter $P_{\mathrm{div}}$ is estimated from newly budded daughter cells originating from young (in terms of RLS) mother cells. If we assume that such a mother cell retains damage, it follows that the corresponding daughter cell will be close to "damage-free" at the point of birth. Therefore the first data point, that is the initial area of these daughter cells, should be $A(t=0) = (1-s)\,\mathrm{P}_{\text{div}}$ where $s$ is the average size proportion estimated from the "jumps" associated with cell division described previously. Thus, the "dimensionless area" is obtained by scaling the data by the value $P_{\text{div}} = \dfrac{A(t=0)}{(1-s)}$.

Morever, the time variable is rendered dimensionless by scaling it by the growth rate $\mu$. The choice of this scaling factor is motivated by the fact that the time variable for the models are rendered dimensionless by using this factor (see section S1.2). The resulting dimensionless time variable is defined as $\tau = \mu \cdot t$, and the implemented value of the growth rate for the model validation is $\mu = 0.5\ \mathrm{h}^{-1}$ for *S.cerevisiae* based on the experiments by Snoep et al.[13]. The above procedure explains how the data is rendered dimensionless, however from the point of view of the experimental design it is important that the ageing phenomena is captured in the data.

As the competing damage accumulation models describe replicative ageing, it is important that the data takes ageing into account. Therefore, we have followed the cell growth of young daughter cells until the first cell division occurs and old mother cells until the last division before cell death occurs. A successful model of replicative ageing should be able to reproduce the growth curves of both a damage-free daughter cell and an old mother cell close to cell death. It is for this reason that we have chosen the following experimental design. By using the "averaging approach" (Fig S2) that will be described next, we generate two separate sets of timeseries data corresponding to an average young daughter cell and an average old mother cell. Thereafter, we say that the model that best can replicate both these two data sets is the one deemed the most realistic.

The four steps of the "averaging"-processing of the data are the following. First, we collect the data of, in this case, three damage-free daughter cells and five old mother cells close to cell death (Fig S2A). Secondly, we extract the growth curves of a single division corresponding to the growth until the first division in the case of the daughter cells and the growth until the last division before cell death in the case of the mother cells (Fig S2B). Thirdly, the area is non-dimensionalised by scaling it by the value $P_{\text{div}}$ (Fig S1) as described previously and the time is scaled so that each cell division occurs after 1 unit of time (Fig S2C). Then, by using interpolation we average the cell area for the different cells and we multiply the the time with the average generation time for all cells in minutes before scaling the time with the growth rate $\mu$ in order to obtain the dimensionless time $\tau$ (Fig S2D). Thus, in the last step the time is "stretched back" by scaling it by both the average generation time and the growth rate $\mu$. The results are the two time series corresponding to an average "damage-free" daughter cell and an average old mother cell close to cell death. The non-dimensionalisation of both the data and the model results in the possibility of quantifying the three modelling parameters $s$, $P_{\text{div}}$ and $Q$ (Tab S1).

**Table S1. Parameters estimated from the data**. The left hand column displays the parameter and the right hand column shows the value of the corresponding parameter.

| **Parameter** | **Value** |
|---|---|
| $s$ | 0.6370 |
| $P_{\text{div}},\ \left[(\mu\mathrm{m})^2\right]$ | 28.8422 |
| $Q$ | 2.5526 |

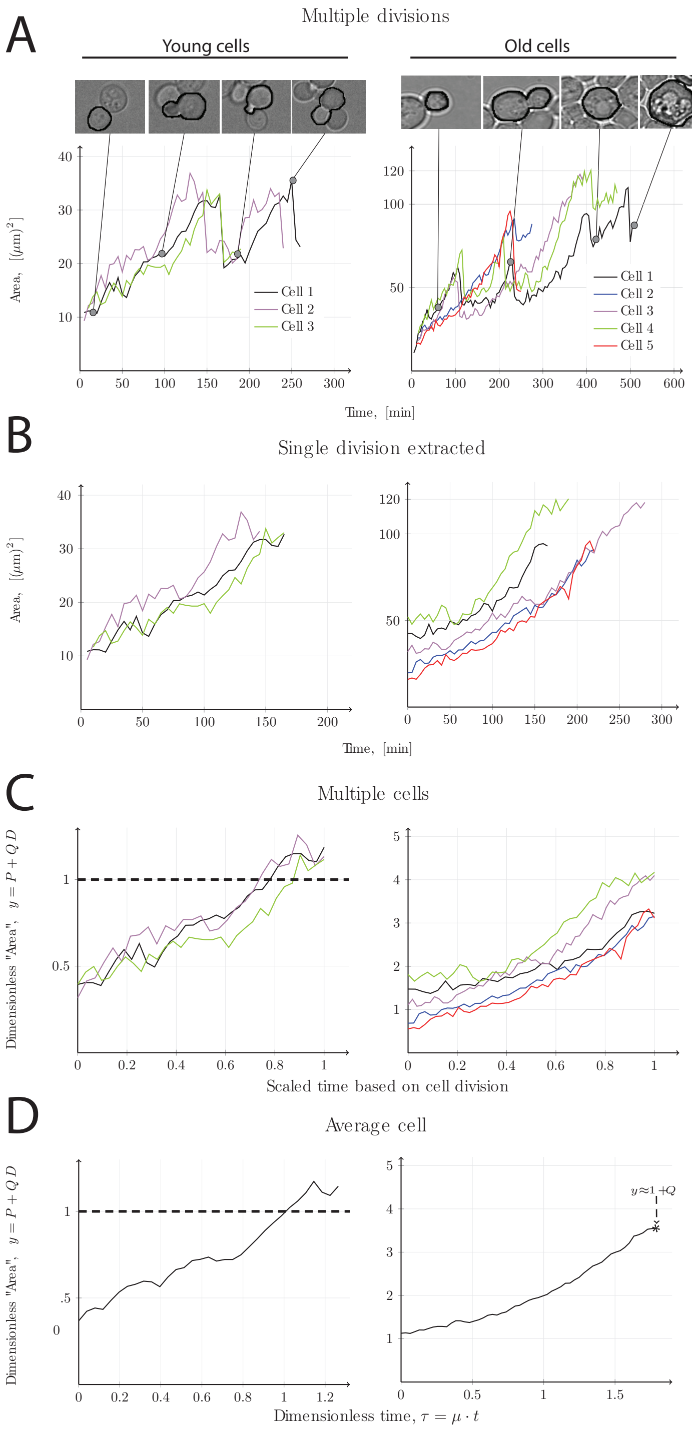

**Figure S2. Procedure for averaging the data**. The left hand column shows the ”damage-free” daughter cells and the right hand column shows the old mother cells. The four rows shows the ”data-averaging” procedure. **(A)** Raw data of multiple cells. **(B)** The extraction of one cell division (the first and last respectively). **(C)** The scaling of data and shrinking of the time so that division occurs after 1 (dimensionless) time unit. **(D)** The average area and the time $\tau$ is stretched back using the growth rate $\mu = 0.5\ \mathrm{h}^{-1}$

Subsequently, the non-dimensionalisation of the competing models, which is compatible the methodology implemented for processing of the data, is presented.

#### S1.2 Non-dimensionalisation of the competing models

The aim is to render the modelling output dimensionless in a similar manner to the non-dimensionalisation of the data. As the modelling output is defined as $\hat{y}(\theta,t) = P(\theta,t) + D(\theta,t)$, this achieved by scaling both the time variable $t$ and the two states $P, D$ appropriately.

The dimensionless time $\tau$ is defined as follows.

$$\tau = \mu\, t \Leftrightarrow t = \frac{\tau}{\mu} \tag{S1}$$

Furthermore, the dimensionless states are the intact and damaged proteins as functions of their respective thresholds $P_{\text{div}}$ and $D_{\text{death}}$ respectively.

$$P \leftarrow \frac{P}{P_{\text{div}}} \quad \& \quad D \leftarrow \frac{D}{D_{\text{death}}} \tag{S2}$$

It is important to emphasise that since $\mu$ is used to introduce the dimensionless time variable $\tau$, it is removed from the presented model during the non-dimensionalisation procedure. Therefore, the growth rate $\mu$ cannot be estimated using the dimensionless model and in order to compare the dimensionless model to the data it is essential that the data is rendered dimensionless in the same way. For this purpose, we have implemented the value $\mu = 0.5$ h$^{-1}$ when the time series data was rendered dimensionless (see section S1.1).

Now, among the previous models there are two that explicitly focus on the bakers yeast *S.cerevisiae*. These models were constructed by Erjavec et al.[8] and Clegg et al.[6]. Using the above non-dimensionalisation procedure, the dimensionless versions of the Erjavec-model (Eq (S3))

$$\begin{aligned} \frac{\mathrm{d}P}{\mathrm{d}\tau} &= \alpha\left(\frac{P}{K_M + P + QD}\right) - k_1 P \\ \frac{\mathrm{d}D}{\mathrm{d}\tau} &= \frac{k_2}{Q}P - k_3 D \end{aligned} \tag{S3}$$

and the Clegg-model (Eq (S4))

$$\begin{aligned} \frac{\mathrm{d}P}{\mathrm{d}\tau} &= (1-\beta)\,P\left(\frac{P}{P + QD}\right) - k_1 P + k_2 D\left(\frac{P}{\beta P + QD}\right) \\ \frac{\mathrm{d}D}{\mathrm{d}\tau} &= \frac{k_1}{Q}P - \frac{\beta}{\mu}D\left(\frac{P}{\beta P + QD}\right) \end{aligned} \tag{S4}$$

are obtained. The differences between the various models are the assumptions underlying the construction of the ODEs which involves the forces cell growth, formation and repair of damage.

In the case of the Erjavec-model (Eq (S3)), the cell growth term is not exponential and the growth rate declines with increasing amounts of damage according to the fraction $\frac{P}{K_M+P+QD}$. Moreover, a fraction of the intact proteins contributes to the formation of damage while the remaining part of the intact proteins is degraded. Lastly, the damage that is repaired does not contribute to the amount of intact proteins but it is degraded instead.

In the case of the Clegg-model (Eq (S4)), the cell growth term is exponential and the growth rate declines as previously with increasing amounts of damage according to the fraction $\frac{P}{P+QD}$. The formation of damage term is the same as in the current model (Eq (S7)). Lastly, a fraction of the damage that is repaired contributes to the amount of intact proteins while the remaining part is degraded and the rate of repair declines with increasing amounts of damage according to the fraction $\frac{P}{\beta P+QD}$.

Using the scaled forms of the variable (Eq (S1)) and the states (Eq (S2)), the three (i.e. the presented model, the model by Erjavec et al.[8] and the model by Clegg et al.[6]) non-dimensionalisation procedures are presented subsequently before the structural identifiability analysis is described.

##### S1.2.1 Non-dimensionalisation of the presented model

The aim is to render the ODE-model (Eq (S5)),

$$\frac{\mathrm{d}P}{\mathrm{d}t} = \mu\,P\left(\frac{g\cdot D_{\text{death}} - D}{D_{\text{death}}}\right) - k_1 P + k_2 D$$

$$\frac{\mathrm{d}D}{\mathrm{d}t} = k_1 P - k_2 D \tag{S5}$$

$$\begin{aligned} P(0) &= f_P\left(s, \text{re}\right), \quad D \in [0, D_{\text{death}}],\; P \in [0, P_{\text{div}}] \\ D(0) &= f_D\left(s, \text{re}\right), \quad t \in \mathbb{R}_+, \quad g \in (1, \infty), \quad k_1, k_2 \in \mathbb{R}_+ \end{aligned}$$

with the following initial conditions (Eq (S6))

$$\begin{aligned} f_P\left(s, \text{re}\right) &= \begin{cases} s\,P_{\text{div}} - \text{re}(1-s)D & \texttt{[M]} \\ (1-s)\,P_{\text{div}} + \text{re}(1-s)D & \texttt{[D]} \end{cases} \\ f_D\left(s, \text{re}\right) &= \begin{cases} \left(s + (1-s)\text{re}\right)D & \texttt{[M]} \\ (1-s)(1-\text{re})D & \texttt{[D]} \end{cases} \end{aligned} \tag{S6}$$

dimensionless. Start by introducing the dimensionless form of the time (Eq (S1)) and the states (Eq (S2)). This results in the fact that the rate parameters becomes scaled by the growth rate $\mu$

$$k_1 \leftarrow \frac{k_1}{\mu} \quad \& \quad k_2 \leftarrow \frac{k_2}{\mu}$$

and the introduction of the *damage resilience* parameter defined as follows.

$$Q = \frac{D_{\text{death}}}{P_{\text{div}}}$$

Now, substituting the time $\tau$ into Eq (S5) yields the following equations

$$\begin{aligned} &\mu\,P_{\text{div}}\frac{\mathrm{d}\left({}^{P}/_{P_{\text{div}}}\right)}{\mathrm{d}\tau} = \\ &\mu\,P_{\text{div}}\left\{\frac{P}{P_{\text{div}}}\left(g - \frac{D}{D_{\text{death}}}\right) - \frac{k_1}{\mu}\frac{P}{P_{\text{div}}} + \frac{k_2}{\mu}\frac{D}{D_{\text{death}}}\left(\frac{D_{\text{death}}}{P_{\text{div}}}\right)\right\} \end{aligned}$$

$$\begin{aligned} &\mu\,D_{\text{death}}\frac{\mathrm{d}\left({}^{D}/_{D_{\text{death}}}\right)}{\mathrm{d}\tau} = \\ &\mu\,D_{\text{death}}\left\{\frac{k_1}{\mu}\frac{P}{P_{\text{div}}}\left(\frac{D_{\text{death}}}{P_{\text{div}}}\right)^{-1} - \frac{k_2}{\mu}\frac{D}{D_{\text{death}}}\right\} \end{aligned}$$

and simplifying the above equations in addition to introducing all scaled components results in the following dimensionless system of ODEs

$$\begin{aligned} \frac{\mathrm{d}P}{\mathrm{d}\tau} &= P(g-D) - k_1 P + k_2 Q D \\ \frac{\mathrm{d}D}{\mathrm{d}\tau} &= \left({}^{k_1}/_{Q}\right) P - k_2 D \end{aligned} \tag{S7}$$

which is the desired result.

□

In a similar fashion, the initial conditions are scaled in the following way

$$f_P\left(s, \mathrm{re}\right) \leftarrow \frac{f_P\left(s, \mathrm{re}\right)}{P_{\mathrm{div}}} \quad \& \quad f_D\left(s, \mathrm{re}\right) \leftarrow \frac{f_D\left(s, \mathrm{re}\right)}{D_{\mathrm{death}}}$$

and substituting these expressions into Eq (S6) yields the following equations.

$$P_{\mathrm{div}} \frac{f_P\left(s, \mathrm{re}\right)}{P_{\mathrm{div}}} = \begin{cases} P_{\mathrm{div}} \left( s - \mathrm{re}(1-s) \dfrac{D}{D_{\mathrm{death}}} \left( \dfrac{D_{\mathrm{death}}}{P_{\mathrm{div}}} \right) \right) & \texttt{[M]} \\ P_{\mathrm{div}} \left( (1-s) + \mathrm{re}(1-s) \dfrac{D}{D_{\mathrm{death}}} \left( \dfrac{D_{\mathrm{death}}}{P_{\mathrm{div}}} \right) \right) & \texttt{[D]} \end{cases}$$

$$D_{\mathrm{death}} \frac{f_D\left(s, \mathrm{re}\right)}{D_{\mathrm{death}}} = \begin{cases} D_{\mathrm{death}} \left( \left(s + (1-s)\mathrm{re}\right) \dfrac{D}{D_{\mathrm{death}}} \right) & \texttt{[M]} \\ D_{\mathrm{death}} \left( (1-s)(1-\mathrm{re}) \dfrac{D}{D_{\mathrm{death}}} \right) & \texttt{[D]} \end{cases}$$

Now, simplifying the above results in addition to using the previous scaling arguments yields

$$\begin{aligned} f_P\left(s, \mathrm{re}\right) &= \begin{cases} s - \mathrm{re}(1-s) Q D & \texttt{[M]} \\ (1-s) + \mathrm{re}(1-s) Q D & \texttt{[D]} \end{cases} \\ f_D\left(s, \mathrm{re}\right) &= \begin{cases} \left(s + (1-s)\mathrm{re}\right) D & \texttt{[M]} \\ (1-s)(1-\mathrm{re}) D & \texttt{[D]} \end{cases} \end{aligned} \tag{S8}$$

which is the desired result.

□

##### S1.2.2 Non-dimensionalisation of the Erjavec model

The continuous part of the original Erjavec model[8] is the following.

$$\begin{aligned} \frac{\mathrm{d}P_{\mathrm{int}}}{\mathrm{d}t} &= \frac{k_1 P_{\mathrm{int}}}{k_s + P_{\mathrm{int}} + P_{\mathrm{dam}}} - \left(k_2 + k_3\right) P_{\mathrm{int}} \\ \frac{\mathrm{d}P_{\mathrm{dam}}}{\mathrm{d}t} &= k_3 P_{\mathrm{int}} - k_4 P_{\mathrm{dam}} \end{aligned}$$

In order to compare the three competing models, the non-dimensionalisation procedure must be conducted in the same way. The reason for this is that the data must also be non-dimensionalised in the same way as the models in order to estimate the rate parameters in the ODEs from the measured data.

To this end, we start off by changing notation of the above equation to obtain the following (Eq (S9)).

$$\begin{aligned}\frac{\mathrm{d}P}{\mathrm{d}t} &= (\alpha\cdot\mu\cdot P_{\text{div}})\,\frac{P}{K_M+P+D}-k_1P\\ \frac{\mathrm{d}D}{\mathrm{d}t} &= k_2P-k_3D\end{aligned}\tag{S9}$$

Above the names of the states are relabelled: $P \leftarrow P_{\text{int}}$ and $D \leftarrow P_{\text{dam}}$. Also the coefficients and rate parameters are relabelled as follows.

$$\begin{aligned}(\alpha\cdot\mu\cdot P_{\text{div}}) &\leftarrow k_1\\ K_M &\leftarrow k_s\\ k_1 &\leftarrow (k_2+k_3)\\ k_2 &\leftarrow k_3\\ k_3 &\leftarrow k_4\end{aligned}$$

Now, the same non-dimensionalisation procedure is conducted on the dynamical system (Eq (S9)) where the states are redefined as follows

$$P \leftarrow \frac{P}{P_{\text{div}}} \quad \& \quad D \leftarrow \frac{D}{D_{\text{death}}}$$

and the time variable is rendered dimensionless as follows.

$$\tau = \mu\, t \Leftrightarrow t = \frac{\tau}{\mu}$$

Introducing the above states, the model is non-dimensionalised as follows.

$$\begin{aligned}&\mu P_{\text{div}}\frac{\mathrm{d}\left({}^{P}/{}_{P_{\text{div}}}\right)}{\mathrm{d}\tau} =\\ &\mu P_{\text{div}}\left\{\alpha\cdot\frac{P_{\text{div}}\left(\frac{P}{P_{\text{div}}}\right)}{P_{\text{div}}\left[\left(\frac{K_M}{P_{\text{div}}}\right)+\left(\frac{P}{P_{\text{div}}}\right)+\left(\frac{D_{\text{death}}}{P_{\text{div}}}\right)\cdot\left(\frac{D}{D_{\text{death}}}\right)\right]}-\frac{k_1}{\mu}\frac{P}{P_{\text{div}}}\right\}\end{aligned}$$

$$\begin{aligned}&\mu D_{\text{death}}\frac{\mathrm{d}\left({}^{D}/{}_{D_{\text{death}}}\right)}{\mathrm{d}\tau} =\\ &\mu D_{\text{death}}\left\{\frac{k_2}{\mu}\left(\frac{P_{\text{div}}}{D_{\text{death}}}\right)\left(\frac{P}{P_{\text{div}}}\right)-\frac{k_3}{\mu}\left(\frac{D}{D_{\text{death}}}\right)\right\}\end{aligned}$$

Now, redefining the states and the time variable as previously described in addition to redefining the parameters as follows

$$
\begin{aligned}
K_M &\leftarrow \frac{K_M}{P_{\text{div}}} \\
k_1 &\leftarrow \frac{k_1}{\mu} \\
k_2 &\leftarrow \frac{k_2}{\mu} \\
k_3 &\leftarrow \frac{k_3}{\mu}
\end{aligned}
$$

results in the following system of ODEs (Eq (S10))

$$
\begin{aligned}
\frac{\mathrm{d}P}{\mathrm{d}\tau} &= \alpha \left( \frac{P}{K_M + P + QD} \right) - k_1 P \\
\frac{\mathrm{d}D}{\mathrm{d}\tau} &= \frac{k_2}{Q} P - k_3 D
\end{aligned}
\tag{S10}
$$

which is the desired result.

□

##### S1.2.3 Non-dimensionalisation of the Clegg model

In a similar fashion as for the Erjavec model, the aim is to render the Clegg model dimensionless[6]. The original model is formulated as follows.

$$
\begin{aligned}
\frac{\mathrm{d}P_{\text{act}}}{\mathrm{d}t} &= (1-\beta)\,\mu P_{\text{act}} \left( \frac{P_{\text{act}}}{P_{\text{act}} + P_{\text{dam}}} \right) - aP_{\text{act}} + \gamma_r \left( \frac{\beta P_{\text{act}} P_{\text{dam}}}{\beta P_{\text{act}} + P_{\text{dam}}} \right) \\
\frac{\mathrm{d}P_{\text{dam}}}{\mathrm{d}t} &= aP_{\text{act}} - \left( \frac{\beta P_{\text{act}} P_{\text{dam}}}{\beta P_{\text{act}} + P_{\text{dam}}} \right)
\end{aligned}
$$

Now, the parameters and states are relabelled as follows.

$$
\begin{aligned}
P &\leftarrow P_{\text{act}} \\
D &\leftarrow P_{\text{dam}} \\
k_1 &\leftarrow a \\
k_2 &\leftarrow \gamma_r \beta
\end{aligned}
$$

which results in the following model (Eq (S11)).

$$
\begin{aligned}
\frac{\mathrm{d}P}{\mathrm{d}t} &= (1-\beta)\,\mu P \left( \frac{P}{P + D} \right) - k_1 P + k_2 D \left( \frac{P}{\beta P + D} \right) \\
\frac{\mathrm{d}D}{\mathrm{d}t} &= k_1 P - \beta D \left( \frac{P}{\beta P + D} \right)
\end{aligned}
\tag{S11}
$$

As previously, the states

$$P \leftarrow \frac{P}{P_{\mathrm{div}}} \quad \& \quad D \leftarrow \frac{D}{D_{\mathrm{death}}}$$

and time variable

$$\tau = \mu\, t \Leftrightarrow t = \frac{\tau}{\mu}$$

are rendered dimensionless. Substituting these dimensionless components into Cleggs model (Eq (S11)) yields the following.

$$\begin{aligned}
&\mu P_{\mathrm{div}} \frac{\mathrm{d}\left({}^{P}/_{P_{\mathrm{div}}}\right)}{\mathrm{d}\tau} = \\
&\mu P_{\mathrm{div}} \Bigg\{ (1-\beta)\, \frac{P}{P_{\mathrm{div}}} \left( \frac{P_{\mathrm{div}}\left(\frac{P}{P_{\mathrm{div}}}\right)}{P_{\mathrm{div}}\left[\left(\frac{P}{P_{\mathrm{div}}}\right) + \left(\frac{D_{\mathrm{death}}}{P_{\mathrm{div}}}\right)\left(\frac{D}{D_{\mathrm{death}}}\right)\right]} \right) - \frac{k_1}{\mu} \frac{P}{P_{\mathrm{div}}} \\
&\quad + \frac{k_2}{\mu} \frac{D_{\mathrm{death}}}{P_{\mathrm{div}}} \frac{D}{D_{\mathrm{death}}} \left( \frac{P_{\mathrm{div}}\left(\frac{P}{P_{\mathrm{div}}}\right)}{P_{\mathrm{div}}\left[\beta\left(\frac{P}{P_{\mathrm{div}}}\right) + \left(\frac{D_{\mathrm{death}}}{P_{\mathrm{div}}}\right)\left(\frac{D}{D_{\mathrm{death}}}\right)\right]} \right) \Bigg\}
\end{aligned}$$

$$\begin{aligned}
&\mu D_{\mathrm{death}} \frac{\mathrm{d}\left({}^{D}/_{D_{\mathrm{death}}}\right)}{\mathrm{d}\tau} = \\
&\mu D_{\mathrm{death}} \Bigg\{ \frac{k_1}{\mu}\left(\frac{P_{\mathrm{div}}}{D_{\mathrm{death}}}\right)\left(\frac{P}{P_{\mathrm{div}}}\right) - \frac{\beta}{\mu}\left(\frac{D}{D_{\mathrm{death}}}\right) \left( \frac{P_{\mathrm{div}}\left(\frac{P}{P_{\mathrm{div}}}\right)}{P_{\mathrm{div}}\left[\beta\left(\frac{P}{P_{\mathrm{div}}}\right) + \left(\frac{D_{\mathrm{death}}}{P_{\mathrm{div}}}\right)\left(\frac{D}{D_{\mathrm{death}}}\right)\right]} \right) \Bigg\}
\end{aligned}$$

Introducing the following parameters

$$\begin{aligned}
k_1 &\leftarrow \frac{k_1}{\mu} \\
k_2 &\leftarrow \frac{k_2}{\mu}
\end{aligned}$$

in addition to the previously defined damage resilience quotient $Q$ yields the following system of ODEs (Eq (S12))

$$\begin{aligned}
\frac{\mathrm{d}P}{\mathrm{d}\tau} &= (1-\beta)\, P \left(\frac{P}{P+QD}\right) - k_1 P + k_2 D \left(\frac{P}{\beta P + QD}\right) \\
\frac{\mathrm{d}D}{\mathrm{d}\tau} &= \frac{k_1}{Q} P - \frac{\beta}{\mu}\, D \left(\frac{P}{\beta P + QD}\right)
\end{aligned} \tag{S12}$$

which is the desired result.

□

#### S1.3 Structural identifiability (SI) of the models reveals that $Q$ cannot be estimated in the model fitting

Although Q can be estimated from the actual data points, it is of interest to see which parameters in the various models that can be estimated from a least square fitting of the models to the times series data of cell area over time. From a theoretical point of view, the formulation of the various ODEs determines which involved parameters that can be estimated from a given output and this can be systematically investigated using what is called a *structural identifiability (SI) analysis.*

For the purpose of conducting such an analysis, the three candidate damage accumulation models have the following structure

$$\begin{aligned}\frac{\mathrm{d}P}{\mathrm{d}\tau} &= f_1\left(P, D, \theta, \tau\right)\\ \frac{\mathrm{d}D}{\mathrm{d}\tau} &= f_2\left(P, D, \theta, \tau\right)\\ P(0) &= f_P\left(s, \mathrm{re}\right)\\ D(0) &= f_D\left(s, \mathrm{re}\right)\\ \hat{y}(\theta, \tau) &= P(\theta, \tau) + Q\, D(\theta, \tau)\end{aligned} \tag{S13}$$

where the functions $f_1$ and $f_2$ govern the dynamics of the system and these functions are determined by the forces cell growth, formation and repair of damage. The initial conditions are determined by the functions $f_P$ and $f_D$ (Eq (S8)) and the vector $\theta$ contains all rate parameters. The structure of the system of ODEs (i.e. the functions $f_1$ and $f_2$) determines which parameters in $\theta$ that can be identified from the output $\hat{y}\left(\theta, \tau\right)$. The parameters that can be identified from the output $\hat{y}\left(\theta, \tau\right)$ for a given model are called *structurally identifiable* and we denote these by $\overset{\sim}{\theta}$. The results of the structural identifiability analysis (Tab S2) of the three competing models have been obtained using the Mathematica software developed by Karlsson et al.[10].

**Table S2. The structural identifiable parameters of the three competing models**. The first column lists the three models (the current, Erjavec et al.[8] and Clegg et al.[6]), the second column lists all parameters $\theta$ in each model and the third column lists the structurally identifiable parameters $\overset{\sim}{\theta}$ from the the output $\hat{y}\,(\theta, \tau)$.

| **Model** | **Parameters** $\theta$ | **Structurally identifiable parameters using the output** $\hat{y}\,(\theta,\tau) = P\,(\theta,\tau) + Q\,D\,(\theta,\tau)$ $\overset{\sim}{\theta}$ |
|---|---|---|
| The presented model (Eq (S7)) | $(k_1, k_2, g, Q)$ | $(k_1, k_2, g)$ |
| Erjavec et al.[8] (Eq (S10)) | $(\alpha, K_M, k_1, k_2, k_3, Q)$ | $(\alpha, K_M, k_1, k_2, k_3)$ |
| Clegg et al.[6](Eq (S12)) | $(\beta, k_1, k_2, Q)$ | $(\beta, k_1, k_2)$ |

Thus, next we estimate the structurally identifiable parameters with the given output $\hat{y}\,(\theta, \tau) = P\,(\theta, \tau) + Q\,D\,(\theta, \tau)$ and where we implement the previously estimated value of $Q$ (Tab S1).

#### S1.4 Model validation shows that the presented model outperforms other similar models but that the parameters are not numerically identifiable

The models are validated by fitting the SI-parameters to the experimental data. The model that best replicates the data of both the young and old average yeast cell (Fig S2) is deemed most accurate. However, even though the optimal parameters in the selected model are structurally identifiable, it is not necessarily the case that these parameters can be identified from the data and then the actual parameters cannot be used for predictions.

For the parameter estimation, start by denoting the non-dimensionalised measured data $y(\tau)$. Further, assume that the data has been measured for $m \in \mathbb{Z}^+$ distinct time points $\tau_i$, $i = 1, 2, \ldots, m$. Then, the underlying assumption of the classical least square parameter estimation procedure is that the error $e$ between the measured data $y$ and the simulated output $\hat{y}$ is normally distributed with zero mean and variances $\sigma_i$.

$$e_i = e\left(\overset{\sim}{\theta}, \tau_i\right) = y(\tau_i) - \hat{y}\left(\overset{\sim}{\theta}, \tau_i\right) \sim \mathcal{N}\,(0, \sigma_i)\,,\; i = 1, 2, \ldots, m \tag{S14}$$

Thus, the idea is to select the structurally identifiable parameters $\overset{\sim}{\theta}$ that minimise the magnitude of the errors $e_i$ for all time points (i.e. $i = 1, 2, \ldots, m$) or equivalently selecting the parameters that minimise the variance $\sigma$. This is equivalent of selecting the parameters that minimise the least square cost function denoted $\text{LS}\left(\overset{\sim}{\theta}\right)$ and accordingly the parameter estimation problem can be formulated as follows.

$$\text{Find} \quad \widetilde{\theta} = \text{argmin LS}\left(\widetilde{\theta}\right) = \text{argmin} \sum_{i=1}^{m}\left(y(\tau_i) - \hat{y}\left(\widetilde{\theta}, \tau_i\right)\right)^2 \tag{S15}$$

When selecting models, it is often necessary to take the complexity of the models into account in addition to the fit. An example of such a criteria for selecting models is the *Akaike Information Criteria* (AIC) and this penalises a large number of parameters [9]. It is defined as follows

$$\text{AIC} = -2\text{log}(L) + 2\,p \tag{S16}$$

where $L$ is the likelihood function and $p$ is the number of parameters. Under the normality assumption (Eq (S14)), the loglikelihood can be written as follows.

$$-\log(L) = \sum_{i=1}^{m}\left[\frac{1}{2}\left(\frac{y(\tau_i) - \hat{y}\left(\widetilde{\theta}, \tau_i\right)}{\sigma_i}\right)^2 + \log\left(\sqrt{2\,\pi}\sigma_i\right)\right] \tag{S17}$$

In our case, we have conducted a classic least square estimation using the built in function ”`lsqnonlin`” in Matlab and in this case only the least square (Eq (S15)) is obtained. Since the error variances $\sigma_i$ (Eq (S14)) often are correlated it can be hard to reconstruct the loglikelihood function (Eq (S17)) from the least square LS. To this end we have assumed a constant variance $s^2$ estimated for the obtained errors $e_i$ as follows.

$$\begin{gathered} s^2 = \frac{1}{m-1}\sum_{i=1}^{m}(e_i - \overline{e})^2 \\ \text{where} \\ \overline{e} = \frac{1}{m}\sum_{i=1}^{m} e_i \end{gathered} \tag{S18}$$

In other words, we assume no correlation between the measurment errors $e_i$ (Eq (S14)) which entails that we assume that $s = \sigma_i \forall i \in \{1, \ldots, m\}$. Thus, we have approximated the loglikelihood (Eq (S17)) as follows

$$\begin{aligned} -\text{log}(L) &= \frac{1}{2\,s^2}\sum_{i=1}^{m}\left(y(\tau_i) - \hat{y}\left(\widetilde{\theta}, \tau_i\right)\right)^2 + m\,\log\left(\sqrt{2\,\pi}s\right) \\ &= \frac{1}{2\,s^2}\,\text{LS}\left(\widetilde{\theta}\right) + m\,\log\left(\sqrt{2\,\pi}s\right) \end{aligned} \tag{S19}$$

and our AIC-values (Eq (S16)) for the three models are calculated using the above approximation (Eq (S19)).

Using the ”least square”-methodology (Eq (S15)), the three rivalling models have been validated (Tab S3, Fig S3). Two major conclusions can be drawn. Firstly, the presented model (Eq (S7)) has a substantially better fit compared to the rivalling models. Therefore, the structure of the model is therefore the most biologically realistic. Secondly, the estimated parameters are not *numerically identifiable (NI)*. The numerical identifiability of the parameters are determined by two factors, namely the relative size of the parameter variance compared to the parameter value itself and the correlation between the various parameters [7]. An unbiased estimator of the parameter variance is called the *coefficient of variation* defined as the quotient of the parameter variance and the parameter value itself, i.e. $c = \frac{\sigma_k}{k}$. In order for the parameter $k$ to be numerically identifiable, the value of $c$ should be close to zero percent. For all three models the coefficients of variation are very large (Tab S3) and therefore the parameter values are not numerically identifiable. In addition, the values of the correlations between the parameters should be close to 0 as well in order for the parameters to be numerically identifiable, which is not the case for any of the three models either.

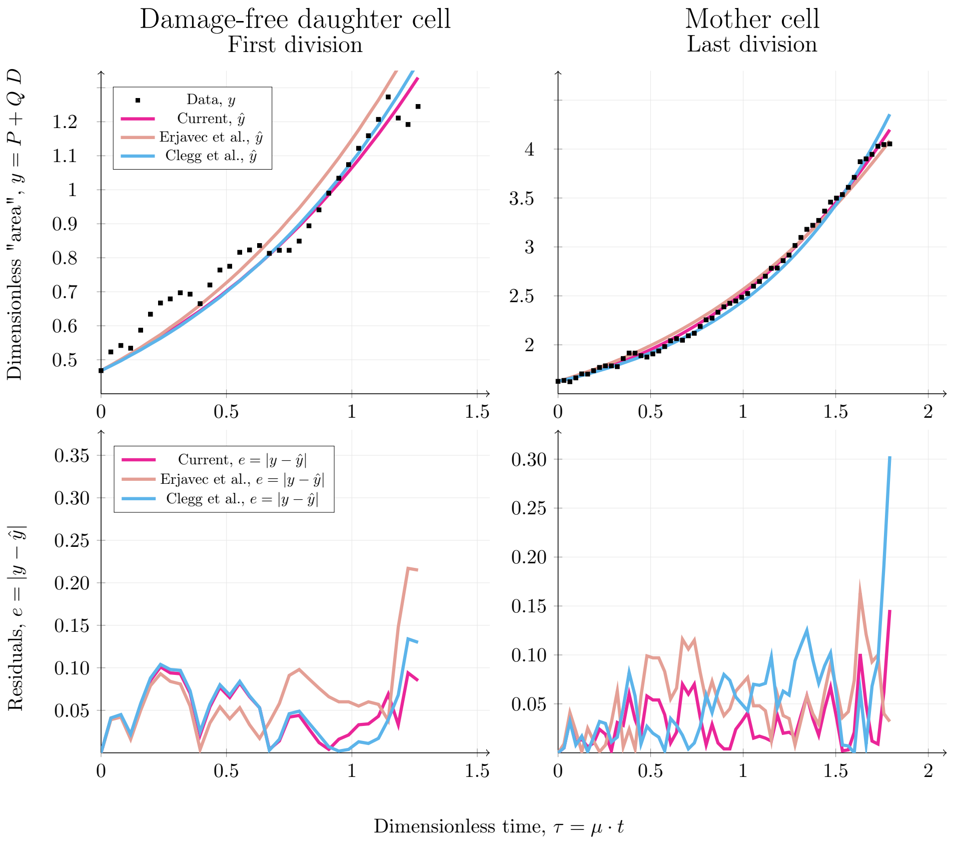

**Figure S3. Validation of the three competing models**. The increase in cell area is plotted against time. The left columns corresponds to a young daughter cell until the first cell division and the right column corresponds to an old mother cell until the last division before cell death. The top row displays the increase in cell area over time while the bottom row shows the evolution of the error $e$ between the measured data $y$ and the model $\hat{y}$. The dashed point show the data points, and the solid curves correspond to the optimal fit of the three competing model. The fit of the three models are: $\left[\text{Magenta: } \left(\text{Model}, r^2\right) = (\text{Current}, 0.20)\right]$,
$\left[\text{Orange: } \left(\text{Model}, r^2\right) = (\text{Erjavec et al.[8]}, 0.46)\right]$ and $\left[\text{Blue: } \left(\text{Model}, r^2\right) = (\text{Clegg et al.[6]}, 0.43)\right]$. For the non-dimensionalisation of the time $\tau$, the growth rate $\mu = 0.5 \; \left[\text{h}^{-1}\right]$ was implemented [13].

**Table S3. Model validation and numerically identifiability**. The five columns are from left to right: The model for which the validation has been conducted, the best fit in terms of the least square (LS), the Akaike Information Criteria (AIC), the optimal parameters obtained from the estimation and the coefficient of variation for each optimal parameter in percent.

| Model | Best Fit LS | AIC | Start guess $\theta_0$ | Lower bound | Upper bound | Optimal Parameters $\theta$ | Coefficient of variation in percent $c = \frac{\sigma_k}{k}$ |
|---|---|---|---|---|---|---|---|
| The presented model (Eq (S7)) $\begin{pmatrix} k_1 \\ k_2 \\ g \end{pmatrix}$ | 0.20 | $-287.67$ | $\begin{pmatrix} 0.50 \\ 0.50 \\ 1.45 \end{pmatrix}$ | $\begin{pmatrix} 0.00 \\ 0.00 \\ 0.90 \end{pmatrix}$ | $\begin{pmatrix} 2.00 \\ 2.00 \\ 2.00 \end{pmatrix}$ | $\begin{pmatrix} 0.15 \\ 0.86 \\ 1.02 \end{pmatrix}$ | $\begin{pmatrix} 7207.5 \\ 486.9 \\ 1982 \end{pmatrix}$ |
| Erjavec et al.[8](Eq (S10)) $\begin{pmatrix} \alpha \\ K_M \\ k_1 \\ k_2 \\ k_3 \end{pmatrix}$ | 0.46 | $-209.75$ | $\begin{pmatrix} 5.45 \\ 5.00 \\ 0.50 \\ 0.50 \\ 0.50 \end{pmatrix}$ | $\begin{pmatrix} 0.9 \\ 0.00 \\ 0.00 \\ 0.00 \\ 0.00 \end{pmatrix}$ | $\begin{pmatrix} 10.00 \\ 10.00 \\ 1.00 \\ 1.00 \\ 1.00 \end{pmatrix}$ | $\begin{pmatrix} 10.00 \\ 8.91 \\ 2.49 \cdot 10^{-13} \\ 6.43 \cdot 10^{-13} \\ 1.99 \cdot 10^{-12} \end{pmatrix}$ | $\begin{pmatrix} 17.78 \\ 19.34 \\ 7.40 \cdot 10^{15} \\ 1.66 \cdot 10^{15} \\ 2.40 \cdot 10^{14} \end{pmatrix}$ |
| Clegg et al.[6](Eq (S12)) $\begin{pmatrix} \beta \\ k_1 \\ k_2 \end{pmatrix}$ | 0.43 | $-219.36$ | $\begin{pmatrix} 0.50 \\ 0.50 \\ 0.50 \end{pmatrix}$ | $\begin{pmatrix} 0.00 \\ 0.00 \\ 0.00 \end{pmatrix}$ | $\begin{pmatrix} 1.00 \\ 1.00 \\ 1.00 \end{pmatrix}$ | $\begin{pmatrix} 0.04 \\ 0.013 \\ 1.00 \end{pmatrix}$ | $\begin{pmatrix} 1.17 \cdot 10^5 \\ 1.37 \cdot 10^5 \\ 829.2 \end{pmatrix}$ |

In addition to the high coefficient of variation, it is also important that the correlation between the parameters are close to zero [7]. The correlations between the parameters for the presented model are the following.

| | $k_1$ | $k_2$ | g |
|---|---|---|---|
| $k_1$ | 1 | - | - |
| $k_2$ | 0.52 | 1 | - |
| $g$ | 0.67 | $-0.27$ | 1 |

The correlations between the parameters for the model by Erjavec et al. [8] are the following.

| | $\alpha$ | $K_M$ | $k_1$ | $k_2$ | $k_3$ |
|---|---|---|---|---|---|
| $\alpha$ | 1 | - | - | - | - |
| $K_M$ | 1.00 | 1 | - | - | - |
| $k_1$ | 1.00 | 1.00 | 1 | - | - |
| $k_2$ | 0.76 | 0.78 | 0.75 | 1 | - |
| $k_3$ | 0.88 | 0.90 | 0.88 | 0.95 | 1 |

The correlations between the parameters for the model by Clegg et al. [8] are the following.

| | $\beta$ | $k_1$ | $k_2$ |
|---|---|---|---|
| $\beta$ | 1 | - | - |
| $k_1$ | 0.97 | 1 | - |
| $k_2$ | 1.00 | 0.98 | 1 |

We next examined the ability of the models to account for replicative ageing by examining if they satisfy the criteria that RLS $\propto Q$ for a fixed set of rate parameters. Thus, by picking parameters giving rise to a finite RLS for all three models, an increase in Q while keeping the remaining parameters fixed should increase the life span, while a decrease in Q should decrease the life span. The simulations generated by the models reveal that the presented model together with the model by Erjavec et al.[8] satisfy this criteria while the model by Clegg et al.[6] does not (Fig S4).

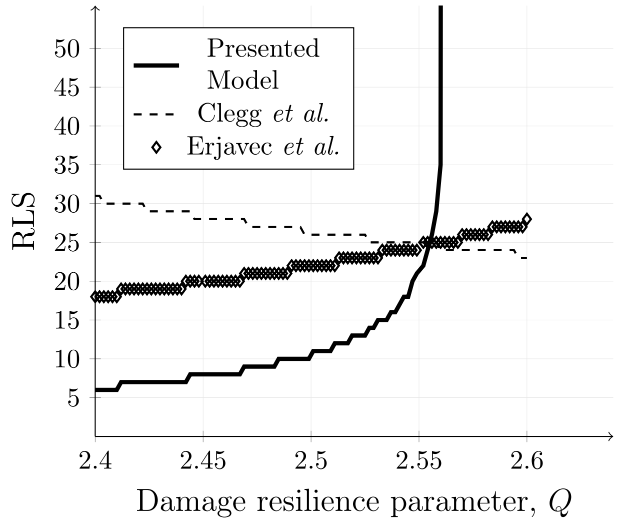

**Figure S4. Replicative life span versus damage resilience.**The replicative life span, RLS, is plotted against the damage resilience, $Q$, for the presented model (black), the model by Erjavec et al[8] (diamonds) and the model by Clegg et al.[6] (dashed black). The specific parameters for the presented model are: $(g, k_1, k_2) = (1.0500, 0.5108, 0.1500)$. The specific parameters for the model by Erjavec et al. are: $(\alpha, K_M, k_1, k_2, k_3) = (2.00, 0.10, 1.00, 0.45, 0.05)$. The specific parameters for the model by Clegg et al. are: $(\beta, k_1, k_2, \mu) = (0.05, 0.32, 0.60, 0.50)$. The overall parameters common for the three models are: $s = 0.6370$, $(P_0, D_0) = (1 - s, 0)$ and where the retention is chosen as a function of the resilience according to re $= \dfrac{1}{Q}\left(\dfrac{2s-1}{1-s}\right)$.

In summary, the following conclusions can be drawn. The current model outperforms the rivalling counterparts in terms of the fit to the experimental data (Tab S3) and it can moreover describe ageing in yeast (Fig S4). However, all of the estimated parameters in all three models are NI indicating that none of the values of the optimal parameters are precise. This implies that there are infinitely many sets of parameters that can give rise to the same fit for all three models. Despite the issues with the identifiability of the parameters, the good fit of the presented model entails that at least parts of the underlying assumptions of the model is true in some sense. Therefore, it is of interest to analyse the mathematical properties of the model at hand.

### S2 Mathematical analysis

Mathematical analysis of the damage accumulation model is conducted to verify its validity. Previously, similar models[1, 8, 12, 3, 4, 6, 11] have answered research questions by means of simulation. However, there are mainly two main problems associated with the implemented methodology in these articles. Firstly, in the construction of these models there has been a lack of motivation as to why a particular model is constructed as it is and whether or not the choice of model parameters is biologically realistic. Secondly, attempts to estimate values of model components and parameters that are very hard or even impossible to measure have been done by searching through the literature. Both these problems can be addressed by investigating the properties of the model at hand by using mathematical analysis. Consequently, the presented damage accumulation model is much more complete as it addresses these two problems by using mathematical analysis as a method in addition to simulations.

As the model is divided into two parts, the discrete (Eq (S8)) and the continuous (Eq (S7)) so is the analysis of the model. The first section concerns the analysis of the discrete part of the models where mathematical relations between the parameters $s$, re and $Q$ are the focus. The second section concerns mainly the parameters $g$, $k_1$ and $k_2$ controlling the dynamics of the continuous part of the model. Here, various constraints on the damage formation rate $k_1$ and the damage repair rate $k_2$ are derived in order to elucidate the allowed sets of parameter that give rise to reasonable dynamics in terms of RLS.

#### S2.1 Analysis of the discrete part of the model reveals the relation between asymmetric division, retention of damage and resilience to damage

##### S2.1.1 The relation between retention, resilience and asymmetry

We wish to derive the following two analytical results

$$0 \leq \text{re} \leq \frac{1}{Q}\left(\frac{2s-1}{1-s}\right)\frac{1}{D} \tag{S20}$$

$$0 \leq s \leq \frac{Q+1}{Q+2} \tag{S21}$$

under the following two assumptions.

1. $P_0$ given by $f_P$ (Eq (S8)) is greater or equal to $P_{0,\min} = (1-s)$
2. $0 \leq \text{re} \leq 1$

For the first result, we know that any mother cell obtains the following proportion of intact proteins after cell division.

$$P_0 = s - \text{re}(1-s)\,Q\,D$$

Since re $\geq 0$ it is the upper bound that must be proven (Eq (S20)). Now, starting from the first assumption listed above yields the following calculations

$$P_0 \geq P_{0,\min}$$

$$\implies s - \mathrm{re}(1-s)\,Q\,D \geq (1-s)$$

$$\begin{aligned}\implies s &\geq (1-s) + \mathrm{re}(1-s)\,Q\,D \\ &= (1-s)\,(1+\mathrm{re}\,Q\,D)\end{aligned}$$

$$\begin{aligned}\implies \frac{s}{(1-s)} &\geq (1+\mathrm{re}\,Q\,D) \\ \iff (1+\mathrm{re}\,Q\,D) &\leq \frac{s}{(1-s)}\end{aligned}$$

$$\begin{aligned}\iff \mathrm{re}\,Q\,D &\leq \frac{s}{(1-s)} - 1 \\ &= \frac{2s-1}{1-s}\end{aligned}$$

$$\iff \mathrm{re} \leq \frac{1}{Q}\left(\frac{2s-1}{1-s}\right)\frac{1}{D}$$

which is the desired result (Eq (S20)).

□

For the second result, we use the second assumption entailing that the maximum amount of damage that the mother cell can retain corresponds to 100%, i.e. $\mathrm{re}_{\max} = 1$. As re $\propto s$ this implies that the following holds.

$$\mathrm{re}_{\max} = \frac{1}{Q}\left(\frac{2s_{\max}-1}{1-s_{\max}}\right)\frac{1}{D}$$

By combining these two findings, the following calculations result in

$$
\begin{aligned}
1 &= \frac{1}{Q}\left(\frac{2s_{\max}-1}{1-s_{\max}}\right)\frac{1}{D} \\
D &= \frac{1}{Q}\left(\frac{2s_{\max}-1}{1-s_{\max}}\right) \\
&\{\text{As } D \leq 1\}
\end{aligned}
$$

$$
\frac{1}{Q}\left(\frac{2s_{\max}-1}{1-s_{\max}}\right) \leq 1
$$

$$
\begin{aligned}
\implies \frac{2s_{\max}-1}{1-s_{\max}} &\leq Q \\
\implies 2s_{\max}-1 &\leq (1-s_{\max})\, Q \\
&= Q - s_{\max}\cdot Q
\end{aligned}
$$

$$
\begin{aligned}
\implies s_{\max}\cdot Q + 2s_{\max} &\leq Q+1 \\
\implies s_{\max}(Q+2) &\leq Q+1
\end{aligned}
$$

$$
\implies s_{\max} \leq \frac{Q+1}{Q+2}
$$

which yields the desired result (Eq (S21)).

□

###### S2.1.2 The increase in generation time as a function of damage resilience

In the article, the effect of the damage resilience parameter on the RLS of individual cells is presented (Fig 1B in the article). The number of divisions is plotted against the time for low ($Q = 2.6$), medium ($Q = 2.8$) and high ($Q = 3.0$) resilence to damage. The length of the ”stairs” in these plots corresponds to the generation time for that division and the generation time increases with the number of divisions. The effect of an increse in resilience to damage on the generation time is shown in the table below (Tab S4).

**Table S4. Generation time for different degrees of resilience to damage**. The four columns are from left to right: number of divisions, the dimensionless generation times for low resilience to damage ($Q = 2.6$), the dimensionless generation times for medium resilience to damage ($Q = 2.8$) and the dimensionless generation times for high resilience to damage ($Q = 3.0$). The other parameters used in the simulations are $g = 1.1$, $k_1 = 0.5$, $k_2 = 0.1$, $s = 0.6370$, $(P_0, D_0) = (1 - s, 0)$ and $\text{re} = 0.2902$.

| **Generation** | **Generation time $\tau_g$** | | |
|---|---|---|---|
| | $Q =$ 2.6 | $Q =$ 2.8 | $Q =$ 3.0 |
| 1 | 1.2340 | 1.1571 | 1.1009 |
| 2 | 1.5330 | 1.3677 | 1.2595 |
| 3 | 1.9299 | 1.6015 | 1.4187 |
| 4 | 2.5485 | 1.8635 | 1.5720 |
| 5 | 6.3693 | 2.1548 | 1.7230 |
| 6 | - | 2.5337 | 1.8684 |
| 7 | - | 3.0919 | 2.0050 |
| 8 | - | 6.9461 | 2.1316 |
| 9 | - | - | 2.2622 |
| 10 | - | - | 2.3934 |
| 11 | - | - | 2.5223 |
| 12 | - | - | 2.6458 |
| 13 | - | - | 2.7611 |
| 14 | - | - | 2.9035 |
| 15 | - | - | 3.0664 |
| 16 | - | - | 3.2471 |
| 17 | - | - | 3.4790 |
| 18 | - | - | 3.8671 |
| 19 | - | - | 7.2382 |
| Mean | 2.7230 | 2.5895 | 2.6034 |

#### S2.2 Analysis of the continuous part of the model results in a theoretical framework describing when replicative ageing occurs in terms of the parameter space

##### S2.2.1 Theoretical framework for the mathematical analysis

In order to pick the parameters $(k_1, k_2)$, two biologically motivated requirements are imposed.

1. Given enough food in the system, the cells should grow and as a consequence of cell growth damage should be accumulated within the cell

2. Every cell within the population has a *finite* replicative life span and should therefore die after a finite number of cell divisions

These biological requirements are now converted into a mathematical description in the following way. The first condition is ensured by using stability analysis and the second by calculating the so called *lethal initial damage threshold*, $D_T$ which determines whether or not a cell will undergo cell death. The idea behind these two methodologies are subsequently presented in chronological order.

The first condition implies that both states $P(\tau)$ and $D(\tau)$ should be *increasing* functions of time which entails calculating the *steady states* of the system. The system, Eq (S7), has two steady states, namely one *trivial* steady state $\mathbf{x}_1^\star = (P_1^\star, D_1^\star) = (0, 0)$ at the origin, and one *non-trivial* steady state at $\mathbf{x}_2^\star = (P_2^\star, D_2^\star) = \left(\frac{k_2 Q}{k_1} g, g\right)$. Thus, the first requirement can be mathematically translated into two parts. Firstly, it is essential to choose the parameters so that the non-trivial steady lies *outside* the square with side length 1 in the phase portrait, that is the point $\mathbf{x}_2^\star$ should be located outside the square $S$ (Eq (S22)).

$$S \coloneqq \{(P, D) \in [0, 1] \times [0, 1]\} \tag{S22}$$

Secondly, the parameters $(k_1, k_2)$ should be chosen based on the *stability* of the fixed points. More precisely, the parameters should be chosen such that the trivial steady state $\mathbf{x}_1^\star$ is *unstable* and the non-trivial steady state $\mathbf{x}_2^\star$ is *stable*. As a consequence, this choice of parameters results in solutions to the system in Eq (S7) that moves *away from* the trivial steady state $\mathbf{x}_1^\star$ and *towards* the non-trivial steady state $\mathbf{x}_2^\star$. In order to illustrate the mentioned dynamics, the proportion of intact proteins $P$ can be plotted against the damage $D$ in a so called *phase portrait* (Fig S5).

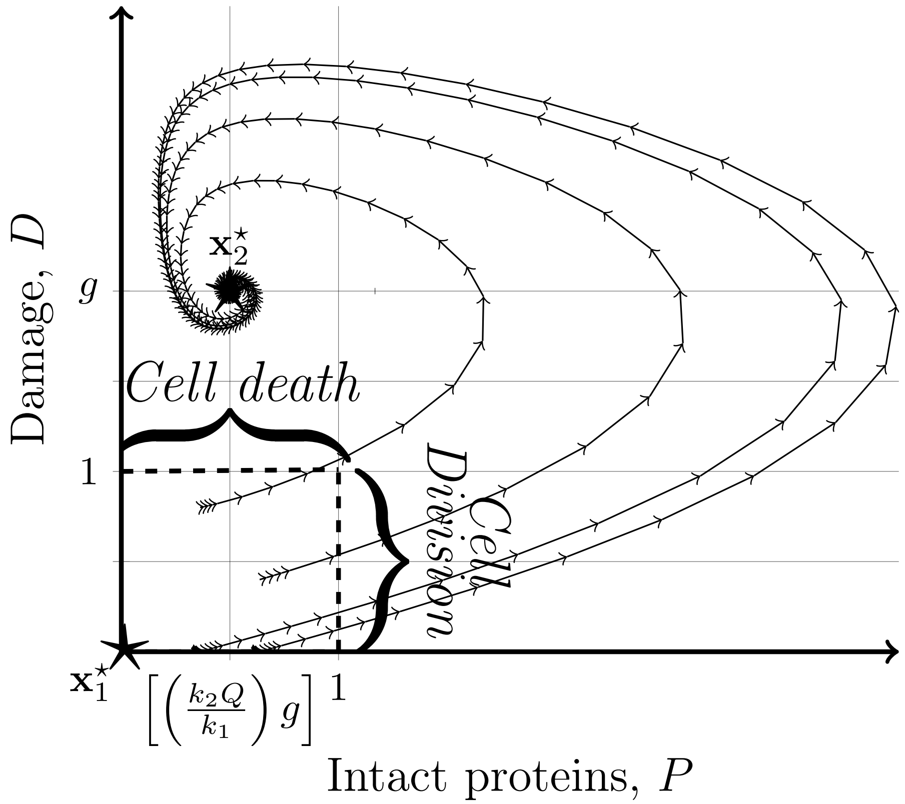

**Figure S5. Phase Portrait of the model**. For each trajectory, the damage $D$ is plotted against the intact proteins $P$ where the time is depicted by the arrows on the trajectories. The trajectories move towards the non-trivial *stable* steady state at $\mathbf{x}_2^\star = \left(\frac{k_2 Q}{k_1} g, g\right)$ and the trajectories move away from the trivial *unstable* steady state at $\mathbf{x}_1^\star = (0, 0)$. In the damage accumulation model, the trajectories are confined to the square $S$ (Eq (S22)) and if they pass through the side where $D = 1$ cell death occurs. Moreover, if the trajectories pass through the side of $S$ where $P = 1$ cell division occurs. The other parameters used in the simulations were: $k_1 = 0.20$, $k_2 = 0.15$, $Q = {}^1/_3$, and $g = 2$.

For the second requirement it is necessary to derive the so called *lethal initial damage threshold* $D_T$. In fact, the mentioned threshold corresponds to a particular trajectory in the phase portrait which determines whether a cell with certain rate parameters will undergo cell death or not. More specifically, if a cell has less damage then $D_T$ at any given time it will undergo at least one more division. Conversely, if the cell in question contains more damage than $D_T$ it will undergo cell death. In order to select parameters that ensure a finite RLS, the values of $D_T$ for both the daughter and mother cells are of interest. For parameter values such that the non-trivial fixed point $x_2^\star$ (Fig S5) is located far away from the cell death side $D = 1$ of the square S (Eq (S22)) this trajectory will be approximately linear and can in this case be approximated using the methodology introduced by Rashidi et al.[12]. More precisely, for these values the trajectory corresponding to the threshold $D_T$ can be obtained by approximating it with numerous lines in the phase portrait. However, for small values of $g$ where $g \approx 1$, the trajectories will be very curved as can be seen in Fig S5. We still do not know how to know how to calculate this trajectory analytically and we ensure the second requirement on the RLS by means of simulations. In other words, given the rate parameters of the model at hand we simulate the RLS by solving the ODEs numerically.

Using the described approach, the aim of the mathematical analysis is to find biologically relevant parameters. Subsquently, the stability analysis is described.

##### S2.2.2 Stability analysis of the damage accumulation model

By using *stability analysis* it is possible to predict the long term dynamical behaviour of a given system. For non-linear systems of ODEs such as the damage accumulation model in this article (Eq (S7)) this goal is achieved in two steps. Firstly, the system is linearised around each steady state and secondly the stability of these linearised systems is determined. For the purpose of predicting the long term dynamics in order to ensure that the first previously posed biological requirement is satisfied, the stability of the formulated damage accumulation model is analysed.

The stability analysis is divided into four steps.

1. Find the two (i.e. the trivial and non-trivial) steady states of the damage accumulation model
2. Derive algebraic expressions for the Jacobian matrix, its determinant and trace
3. Find constraints on the model parameters such that the non-trivial steady state is *stable*
4. Find constraints on the model parameters such that the trivial steady state is *unstable*

The stability analysis begins with finding the steady states to the damage accumulation model (Eq (S7)). Using the notation $(P^\star, D^\star)$, the steady states of interest satisfy the following equations.

$$0 = P^\star\left(g - D^\star\right) - k_1 P^\star + k_2 Q D^\star \tag{S23}$$

$$0 = \frac{k_1}{Q} P^\star - k_2 D^\star \tag{S24}$$

Using Eq (S24) the solution satisfies

$$D^\star = \frac{k_1}{k_2 Q} P^\star \tag{S25}$$

and substituting Eq (S25) into (S23) yields the following calculation.

$$\begin{aligned}
0 &= P^\star\left(g - D^\star\right) - k_1 P^\star + k_2 Q D^\star \\
\implies 0 &= P^\star\left(g - \frac{k_1}{k_2 Q}P^\star\right) - k_1 P^\star + k_2 Q \frac{k_1}{k_2 Q} P^\star \\
\implies 0 &= P^\star\left(\frac{k_2 Q}{k_1}\frac{k_1}{k_2 Q} g - \frac{k_1}{k_2 Q}P^\star\right) - k_1 P^\star + k_1 P^\star \\
\implies 0 &= \frac{k_1}{k_2 Q} P^\star\left(\frac{k_2 Q}{k_1} g - P^\star\right) \\
\implies 0 &= P^\star\left(\frac{k_2 Q}{k_1} g - P^\star\right)
\end{aligned} \tag{S26}$$

Now, Eq (S26) entails that there are two steady states. Combining Eq (S26) and (S25) yields the following two equations

$$P_1^\star = 0 \implies D_1^\star = \frac{k_1}{k_2 Q} P_1^\star = 0 \implies \boxed{\mathbf{x}_1^\star = (P_1^\star, D_1^\star) = (0, 0)} \tag{S27}$$

$$P_2^\star = \frac{k_2 Q}{k_1} g \implies D_2^\star = \frac{k_1}{k_2 Q} P_2^\star = g \implies \boxed{\mathbf{x}_2^\star = (P_2^\star, D_2^\star) = \left(\frac{k_2 Q}{k_1} g, g\right)} \tag{S28}$$

Thus, the damage accumulation model, Eq (S7), has two steady states: a trivial steady state $\mathbf{x}_1^\star$ (Eq (S27)) at the origin and a non-trivial steady state $\mathbf{x}_2^\star$ (Eq (S28)).

□

Next, a general expression for the Jacobian matrix is derived. Using the definition of the Jacobian matrix on the damage accumulation model (Eq (S7)) yields the following result.

$$J\left(\mathbf{x}^\star\right) = J\left(P^\star, D^\star\right) = \begin{pmatrix} (g - D^\star) - k_1 & k_2 Q - P^\star \\ {}^{k_1}/_{Q} & -k_2 \end{pmatrix} \tag{S29}$$

Calculating the trace and determinant of the Jacobian matrix (Eq (S29)) results in the following two equations.

$$\mathrm{Tr}\left[J\left(\mathbf{x}^\star\right)\right] = \mathrm{Tr}\left[J\left(P^\star, D^\star\right)\right] = \left(g - D^\star\right) - \left(k_1 + k_2\right) \tag{S30}$$

$$\mathrm{Det}\left[J\left(\mathbf{x}^\star\right)\right] = \mathrm{Det}\left[J\left(P^\star, D^\star\right)\right] = \left(\frac{k_1 P^\star}{Q} - k_2\left(g - D^\star\right)\right) \tag{S31}$$

Using the formulas for the trace and determinant (Eq (S30) and (S31)) it is possible to determine the stability of the trivial (Eq (S27)) and non-trivial (Eq (S28)) steady state respectively.

□

Next, we would like to find parameters such that the non-trivial steady state is stable. By plugging in the non-trivial steady state, Eq (S28), into Eq (S30) and (S31) in order to obtain expressions for the trace and determinant yields the following.

$$\mathrm{Tr}\left[J\left(\mathbf{x}_2^\star\right)\right] = -\left(k_1 + k_2\right)$$
$$\mathrm{Det}\left[J\left(\mathbf{x}_2^\star\right)\right] = g\,k_2$$

According to the definition of stability, the non-trivial steady state (Eq (S28)) is stable for *positive* rate parameters, i.e. $k_1, k_2 > 0$.

□

Lastly, we would like to find parameters such that the trivial steady state is unstable. Again plugging in the trivial steady state, Eq (S27), into Eq (S30) and (S31) the following expressions are obtained.

$$\mathrm{Tr}\left[J\left(\mathbf{x}_1^\star\right)\right] = g - \left(k_1 + k_2\right)$$
$$\mathrm{Det}\left[J\left(\mathbf{x}_1^\star\right)\right] = -g\,k_2$$

Given these expression, the aim is to find parameters such that trajectories move away from the trivial steady state $\mathbf{x}_1^\star$. Thus, in order to ensure instability we could choose parameters such that the trace of the Jacobian at the trivial steady state is positive. This is exactly what we call the *starvation constraint* in the triangle of ageing which is formulated as follows.

$$g > k_1 + k_2 \tag{S32}$$

The reason why it is called the ''starvation constraint" is due to the fact that it connects the uptake of food, i.e. $g$, to the rate of formation of damage and repair, i.e. $k_1$ and $k_2$. For these parameters the trivial steady state will definitely be *unstable*.

However, potential problems might occur since the determinant $\mathrm{Det}\left[J\left(\mathbf{x}_1^\star\right)\right]$ is always *negative*. The negativity of the term is caused by the fact that the uptake of food is positive, i.e. $g > 0$, as a consequence of Eq (S32) as well as the repair rate, i.e. $k_2 > 0$. Consequently, for parameters satisfying the starvation constraint the trace will be positive, $\text{Tr}\left[J\left(\mathbf{x}_1^\star\right)\right] > 0$, and the determinant will be negative, $\text{Det}\left[J\left(\mathbf{x}_1^\star\right)\right] < 0$. These parameters will therefore give rise to one positive and one negative eigenvalue and consequently the trivial steady state $\mathbf{x}_1^\star$ is characterised as a *saddle point*. This implies that solutions will be drawn towards the trivial steady state along the eigenvector corresponding to the negative eigenvalue and move away from the trivial steady state along the other eigenvector corresponding to the positive eigenvalue. Since the damage accumulation model is restricted to the square $S$ (Eq (S22)) it is important that the eigenvector corresponding to the negative eigenvalue does *not* influence the dynamics within $S$. Therefore, it is necessary to ensure that the ''stable" eigenvector is not located in either the first or third quadrant.

More precisely, assume that the negative eigenvalue of $J\left(\mathbf{x}_1^\star\right) = J\left(0,0\right)$ is called $\lambda$. Define its associated eigenvector $\mathbf{v}$.

$$\mathbf{v} = \begin{pmatrix} v_1 \\ v_2 \end{pmatrix}$$

Then the task at hand can be formulated as follows. In order to ensure that trajectories move away from $\mathbf{x}_1^\star$ (Eq (S27)) and towards $\mathbf{x}_2^\star$ (Eq (S28)) *within* the square $S$ (Eq (S22)) the sign of the first, i.e. $v_1$, and second, i.e. $v_2$, element of the eigenvector $\mathbf{v}$ must have *opposite* signs. Since it is always possible to scale the eigenvector $\mathbf{v}$, we can set $v_1 = 1$ without loss of generality. Provided this notation, we need to prove that the other element of $\mathbf{v}$ is always negative, i.e. $v_2 < 0$. By definition, the eigenvector $\mathbf{v}$ satisfies the matrix equation

$$J\left(0,0\right) \cdot \mathbf{v} = \lambda \mathbf{v}$$

which can be written as follows.

$$\begin{pmatrix} g - k_1 & k_2 Q \\ {}^{k_1}/_{Q} & -k_2 \end{pmatrix} \cdot \begin{pmatrix} v_1 \\ v_2 \end{pmatrix} = \lambda \begin{pmatrix} v_1 \\ v_2 \end{pmatrix}$$

The above matrix equation is equivalent to the following two equations.

$$\begin{aligned} \left(g - k_1\right) v_1 + k_2 Q v_2 &= \lambda v_1 \\ \frac{k_1}{Q} v_1 - k_2 v_2 &= \lambda v_2 \end{aligned}$$

Substituting, the value $v_1 = 1$ into the upper of the two equations and solving for $v_2$ yields the following result.

$$v_2 = \frac{\lambda - \left(g - k_1\right)}{k_2 Q}$$

The eigenvector of interest can thus be written as a scalar multiple of the following vector.

$$\mathbf{v} = \begin{pmatrix} 1 \\ \\ \dfrac{\lambda - \left(g - k_1\right)}{k_2 Q} \end{pmatrix} \quad \text{where} \quad \lambda < 0 \tag{S33}$$

As a result of the starvation constraint (Eq (S32)) the term $\left(g - k_1\right)$ must be positive. Since the eigenvalue is negative it follows that the numerator in the element $v_2$ is negative and the denominator is positive. Accordingly the second element of the eigenvector associated with the negative eigenvalue

$\lambda$ is negative, i.e. $v_2 < 0$, and thus the eigenvector $\mathbf{v}$ (Eq (S33)) lies in the fourth quadrant. Thus, for positive parameters which satisfies the starvation constraint the trajectories will move away from the trivial steady state $\mathbf{x}_1^\star$ and towards the non-trivial steady state $\mathbf{x}_2^\star$ within the square $S$ (Eq (S22)).

$\square$

### S3 Numerical implementations

The numerical implementation underlying the results from the simulations yields four key outcomes. Firstly, the RLS of an individual cell is simulated. Secondly, the ageing region of the parameter space is generated. Thirdly, the comparison between the life prolonging strategies is conducted. Fourthly, the selection of parameters by fitting the model to data (see subsection S1.1) is conducted. The scripts which generated the results have been written in Matlab and are available at request. However, in this section the code is not attached but instead the ideas behind the numerical implementations are presented in terms of pseudocode algorithms.

#### S3.1 Simulate the RLS of a single cell

Imagine that we have a differential equation of the following type.

$$\frac{\mathrm{d}\mathbf{x}}{\mathrm{d}t} = f\left(t, \mathbf{x}\right)$$

For stiff problems for which we have a large eigenvalue in terms of its absolute value, the dynamics initially changes quickly and later it stabilises. In these cases it is appropriate to use an adaptive method such as *Runge-Kutta Fehlberg 45, RK45* method[5]. These types of methods are based on a Taylor expansion of the function $f$ where we step forward in time given a discretisation of the time line. For example a first order approximation with a step size $\Delta t$ proceeds as follows

$$\mathbf{x}_1(t_{k+1}) = \mathbf{x}_1(t_k) + \Delta t \cdot f\left(t_k, \mathbf{x}_1(t_k)\right)$$

where $t_{k+1} = t_k + \Delta t$ and $\Delta t$ is rather small in order to get a stable algorithm for stiff problems. In order to get a faster solution algorithm it is possible to derive a fourth $\mathbf{x}_4$ and a fifth $\mathbf{x}_5$ order approximation where the error is given by $e = \|\mathbf{x}_5 - \mathbf{x}_4\|$. Subsequently the step size is adapted as long as the error $e$ is small enough, and this is the idea behind the RK45 method. In Matlab the solver "ode45" uses this algorithm which given a time interval $[t_0, t_{\max}]$ and initial conditions $\mathbf{x}_0 = \mathbf{x}(t_0)$ as inputs the solver returns a vector with time points $\vec{t}$ and corresponding solutions $\vec{\mathbf{x}}$ at these time points as outputs.

For our model, we would provide the maximum time $\tau_{\max}$ and the initial conditions $(P_0, D_0)$ as inputs and then the solver would return the three vectors $\vec{\tau}, \vec{P}$ & $\vec{D}$. Assume that the number of time steps to reach $\tau_{\max}$ which is the length of these vectors is $m$. Then provided that the $\tau_{\max}$ is large enough there should be an index $i \in \{1, \ldots, m\}$ for which the $i^{\text{th}}$ element of the vector $\vec{P}$, denote this $\vec{P}_i$, satisfies the following

$$\vec{P}_i \leq 1 \leq \vec{P}_{i+1}$$

meaning that cell division occurs between these entries in time. Then using linear interpolation we can calculate the step length

$$h = \frac{1 - \vec{P}_i}{\vec{P}_{i+1} - \vec{P}_i}$$

and then we would get the generation time $\hat{\tau}$, the intact proteins at division $P$ and the damage at division $D$ according to the following assignments.

$$\begin{aligned} P &\leftarrow 1 \\ D &\leftarrow \vec{D}_i + h \cdot \left(\vec{D}_{i+1} - \vec{D}_i\right) \\ \hat{\tau} &\leftarrow \vec{\tau}_i + h \cdot \left(\vec{\tau}_{i+1} - \vec{\tau}_i\right) \end{aligned}$$

By plugging in $D$, the size proportion $s$ and the retention value re into the functions $f_P$ and $f_D$ (Eq (S8)) the initial conditions for the next generation is obtained. This process can be continued until

$D$ reaches the value 1 before $P$ does implying that cell death occurs. We present a flowchart of the above scheme below (Alg 1).

**Algorithm 1** Simulation of the RLS for a single cell

**Output:**
  Replicative Life Span: RLS & a vector of generation times $\vec{\hat{\tau}}$
**Input:**
  $\tau_{\max}$, $(P_0, D_0)$, $S \leftarrow 1$ $s \leftarrow \dfrac{3}{4}$, re, $K_S$, $Q$, $g$ $k_1$, $k_2$, $\mu(S) \leftarrow \dfrac{S}{S + K_S}$
**Initialise:**
  Initiate the RLS, i.e. RLS $\leftarrow 0$
*Solve ODEs*: Solve the system of ODEs until $\tau_{\max}$ is reached with the initial conditions $(P_0, D_0)$
**if** $P \geq 1$ for some time $\tau_i$ **then**
  Interpolate P to 1, $\tau_i$ to the generation time $\hat{\tau}$ and $D$ to its corresponding value
**end if**
**if** $D \geq 1$ for some time $\tau_i$ and $P < 1 \forall \tau_i$ **then**
  Clonal senescence has occurred$\Longrightarrow$Terminate
**end if**
Define a help index, $i \leftarrow 1$, and the maximum number of iterations, maxIter $\leftarrow 600$
**while** $D < 1$ & $i <$ maxIter **do**
  Increment the number of generations: RLS $\leftarrow$ RLS $+ 1$
  Save the generation time $\hat{\tau}$ in the generation time vector $\vec{\hat{\tau}}$, that is $\vec{\hat{\tau}}(i) \leftarrow \hat{\tau}$
  *Reinitialisation*: Reinitialise the intact proteins and damage, i.e. reassign values to $(P_0, D_0)$, using the functions $f_P$ and $f_D$, Eq (S8), with $P$, $D$, $Q$, $s$ and re as inputs
  *Solve ODEs*: Solve the system of ODEs until $\tau_{\max}$ is reached with the initial conditions $(P_0, D_0)$
  **if** $P \geq 1$ for some time $\tau_i$ **then**
    Interpolate P to 1, $\tau_i$ to the generation time $\hat{\tau}$ and $D$ to its corresponding value
    Increment the help index: $i \leftarrow i + 1$
  **end if**
  **if** $D \geq 1$ for some time $\tau_i$ and $P < 1 \forall \tau_i$ **then**
    Cell death has occurred$\Longrightarrow$Terminate
  **end if**
**end while**
**return** RLS and $\vec{\hat{\tau}}$

#### S3.2 Generating the replicative ageing region of the parameter space

The ageing are of the parameter space is generated by simulating the RLS of individual cells (Alg 1) for different parameter values. Given the various model parameters it is possible to create two vectors with equidistant values of the rates $k_1$ and $k_2$ where the number of elements is predetermined. Then by looping through these vectors it is possible to save the parameter pairs $(k_1, k_2)$ that satisfies all constraints with their corresponding RLS. The algorithm for generating the ageing region of the parameter space is presented below (Alg 2).

**Algorithm 2** Generating the ageing region of the parameter space

**Output:**
Two lists of rate parameters $\vec{k}_1$ & $\vec{k}_2$ and a list of the corresponding RLSs, $\overrightarrow{\mathrm{RLS}}$

**Input:**
$S \leftarrow 1$, $s \leftarrow \dfrac{3}{4}$, re, $K_S$, $Q$, $g$, $\mu(S) \leftarrow \dfrac{S}{S+K_S}$

**Initialise:**
Define the number of parameters that is to be included in the triangle, e.g. $n \leftarrow 120$. Allocate two mesh vectors with that size, i.e. $\mathbf{k}_1 \leftarrow \mathbf{0} \in \mathbb{R}^{n+1}$ & $\mathbf{k}_2 \leftarrow \mathbf{0} \in \mathbb{R}^{n+1}$. Set two minimum, e.g. $k_{1,\text{min}} \leftarrow 0.001$ & $k_{2,\text{min}} \leftarrow 0.001$, and maximum , e.g. $k_{1,\text{max}} \leftarrow \mu(S)\, g$ & $k_{2,\text{max}} \leftarrow \mu(S)\, g$, values. Define a mesh size $h_j \leftarrow \left(\dfrac{k_{j,\text{max}} - k_{j,\text{min}}}{n}\right)$, $j \in \{1,2\}$ and loop through the two respective meshes where each individual element is assigned the following value $\mathbf{k}_j(i) = k_{j,\text{min}} + (i-1)\cdot h_j,\ i \in \{1,\ldots,n+1\}, j \in \{1,2\}$

**for** $i \in \{1,\ldots,n+1\}$ **do**
  $k_1 \leftarrow \mathbf{k}_1(i)$
  **for** $j \in \{1,\ldots,n+1\}$ **do**
    $k_2 \leftarrow \mathbf{k}_2(j)$
    **if** $(g > k_1 + k_2)$ **then**
      Simulate the RLS using Alg 1
      **if** $0 < \mathrm{RLS} < \infty$ **then**
        Save $k_1$, $k_2$ and RLS in the vectors $\vec{k}_1$, $\vec{k}_2$ and $\overrightarrow{\mathrm{RLS}}$ respectively
      **end if**
    **end if**
  **end for**
**end for**
**return** $\vec{k}_1$, $\vec{k}_2$ and $\overrightarrow{\mathrm{RLS}}$

#### S3.3 Comparing life prolonging strategies

The two strategies for prolonging the RLS are compared by using the two previous algorithms. The aim is to compare the efficiency of a decrease in the damage formation rate and an increase damage repair rate with respect to prolonging the RLS. By generating a set of rate parameters $(k_1, k_2)$ within the triangle of ageing (Alg 2) it is possible to loop over the triangle of ageing and simulate the RLS for a cell with these parameters (Alg 1). Subsequently, the increase in RLS by decreasing the damage formation rate $\Delta\mathrm{RLS}_{k_1}$ is compared to the increase in RLS by increasing the repair rate $\Delta\mathrm{RLS}_{k_2}$ in order to elucidate the most effective strategy. The algorithm for comparing the life prolonging strategies is presented below (Alg 3).

**Algorithm 3** Comparing life prolonging strategies

**Output:**
Three files containing parameter pairs called ‘’file_$k_1$”, ‘’file_$k_2$” and ‘’file_neither”. Also three proportions of points $p_{k_1}$, $p_{k_2}$ and $p_{\text{neither}}$ are returned
**Input:**
$\tau_{\max}$, $(P_0, D_0)$, $S \leftarrow 1$ $s \leftarrow \frac{3}{4}$, re, $K_S$, $Q$, $g$ $k_1$, $k_2$, $\mu(S) \leftarrow \frac{S}{S+K_S}$
**Initialise:**
Define the number of elements in the triangle of ageing, e.g. $n \leftarrow 500$.
Generate a set of rate parameters $(\mathbf{k}_1, \mathbf{k}_2)$ within the triangle of ageing (Alg 2) and assume that the number of parameter pairs within the triangle is $m$, that is $\mathbf{k}_1, \mathbf{k}_2 \in \mathbb{R}^m$.
Allocate memory for a vector with replicative life spans, $\mathbf{RLS} \in \mathbb{R}^m$
**for** $i \in \{1, \ldots, m\}$ **do**
  Define rate parameters, $k_1 \leftarrow \mathbf{k}_1(i)$ and $k_2 \leftarrow \mathbf{k}_2(i)$
  Simulate the RLS with the given rate parameters $(k_1, k_2)$ (Alg 1) , denote this RLS $(k_1, k_2)$.
  Define $\Delta_{k_1} = (0.05 \cdot k_1)$ and simulate the RLS with the given rate parameters $(k_1 - \Delta_{k_1}, k_2)$ (Alg 1) , denote this RLS $(k_1 - \Delta_{k_1}, k_2)$.
  Define $\Delta_{k_2} = (0.05 \cdot k_2)$ and simulate the RLS with the given rate parameters $(k_1, k_2 + \Delta_{k_2})$ (Alg 1) , denote this RLS $(k_1, k_2 + \Delta_{k_2})$.
  **if** $[\text{RLS}\,(k_1 - \Delta_{k_1}, k_2) - \text{RLS}\,(k_1, k_2)] > [\text{RLS}\,(k_1, k_2 + \Delta_{k_2}) - \text{RLS}\,(k_1, k_2)]$ **then**
    $p_{k_1} \leftarrow (p_{k_1} + 1)$
  **end if**
  **if** $[\text{RLS}\,(k_1, k_2 + \Delta_{k_2}) - \text{RLS}\,(k_1, k_2)] > [\text{RLS}\,(k_1 - \Delta_{k_1}, k_2) - \text{RLS}\,(k_1, k_2)]$ **then**
    $p_{k_2} \leftarrow (p_{k_2} + 1)$
  **end if**
  **if** $[\text{RLS}\,(k_1, k_2 + \Delta_{k_2}) - \text{RLS}\,(k_1, k_2)] = [\text{RLS}\,(k_1 - \Delta_{k_1}, k_2) - \text{RLS}\,(k_1, k_2)]$ **then**
    $p_{\text{neither}} \leftarrow (p_{\text{neither}} + 1)$
  **end if**
**end for**
$p_{k_1} \leftarrow ({}^{p_{k_1}}/{}_{m})$
$p_{k_2} \leftarrow ({}^{p_{k_2}}/{}_{m})$
$p_{\text{neither}} \leftarrow ({}^{p_{\text{neither}}}/{}_{m})$
**return** The three files ‘’file_$k_1$”, ‘’file_$k_2$” and ‘’file_neither”, and the three proportions $p_{k_1}$, $p_{k_2}$ and $p_{\text{neither}}$

#### S3.4 Parameter estimation and numerical identifiability analysis using the experimental data

For the parameter estimation used in the model validation (subsection S1.4), there are three functions in Matlab that are required. Firstly, an ODE-solver such as `ode45` is required in order to simulate the model and regenerate the simulated output $\hat{y}(\theta)$ with the given parameters $\theta$. Secondly, an interpolation function is needed in order to interpolate the simulated output to the measured time points. Thirdly, an optimisation function such as `lsqnonlin` is required that iteratively updates the parameter vector $\theta$ until the termination criteria is met.

For the validation step, the models were fitted to the two separate time series of cell growth for an average daughter cell and an average mother cell (Fig S2D). In both cases, the initial conditions for the two time series were different. Despite this fact, the initial conditions of the simulated output were set to interpolate the data at the starting point of the two respective time series, however the initial conditions for the two states of the model were different for the two time series. In the case of the young daugther cell, the initial conditions were set to

$$(P_0, D_0) = (y_1(\tau = 0), 0) = (0.3680, 0.00)$$

were $y_1(\tau = 0)$ is the first data point of the time series corresponding to the dimensionless area of the average daughter. Above, the initial conditions build on the assumption that the daughter cell is born with no damage.

For the time series correponding to the old mother cell, we assume that the mother cell has accumulated damage. If we denote the first data point of the second time series by $y_2(\tau = 0)$, according to the initial conditions, the damage before division is the following.

$$D^\star = \frac{1}{Q}\left(\frac{y_2(\tau = 0)}{s} - 1\right) = 0.3012$$

Using the above notation, the initial conditions (Eq (S8)) are the set to following.

$$(P_0, D_0) = (s - \text{re}(1 - s)QD^\star, (s + (1 - s)\text{re})\, D^\star) = (0.5545, 0.2241)$$

With the above initial conditions for the two respective time series, the results from the model validation part have been obtained (Tab S3).

From the solver `lsqnonlin`, it is also possible to conduct the identifiability analysis. This procedure relies on the Jacobian matrix which contains the time series of the derivatives of the model output with respect to the model parameters. In our case, we have the following output as an approximation of the "dimensionless area"

$$\hat{y}(\theta, \tau) = P(\theta, \tau) + Q\, D(\theta, \tau)$$

and in the case of the presented model (Eq (S7)) the following parameters are estimated from the data.

$$\theta = \begin{pmatrix} g \\ k_1 \\ k_2 \end{pmatrix}$$

Then, the Jacobian matrix is given by

$$J = \begin{pmatrix} \frac{\partial \hat{y}}{\partial g} & \frac{\partial \hat{y}}{\partial k_1} & \frac{\partial \hat{y}}{\partial k_2} \end{pmatrix} \tag{S34}$$

where the columns consist of the so called *sensitivities* [2]. The sensitivities are crucial for the local algorithms that most numerical optimisers use in their iterative search for a local optima, and by using these it is possible to conduct the numerical identifiability analysis. Assume that the estimated variance $s^2$ of the measured data is given by

$$s^2 = \frac{1}{m-1}\sum_{i=1}^{m}\left(y(t = t_i) - \overline{y}\right)^2$$

where $m$ is the number of data points and $\overline{y}$ is the average dimensionless area. In this case, we use the Jacobian matrix (Eq (S34)) to calculate the *Fisher Information matrix* as follows [7].

$$F = \frac{1}{s^2}\left(J^T \cdot J\right) \tag{S35}$$

The inverse of the Fisher information matrix, $F^{-1}$, constitutes a lower bound of the covariance matrix $\text{COV}(\theta)$ and hence we approximate the covariance matrix accordingly.

$$\text{COV}(\theta) \approx F^{-1}$$

In Matlab, it is possible to obtain the correlation matrix by using the function `corrcov` on the above matrix. The coefficient of variation for each parameter is obtained by dividing the corresponding diagonal element of the covariance matrix with the actual parameter values. For further reading see Chapter 12 in the book by DiStefano [7].

7. Joseph DiStefano III. *Dynamic systems biology modeling and simulation*. Academic Press, 2015.

8. Nika Erjavec, Marija Cvijovic, Edda Klipp, and Thomas Nyström. Selective benefits of damage partitioning in unicellular systems and its effects on aging. *Proceedings of the National Academy of Sciences*, 105(48):18764–18769, 2008.

9. EP George Box and M Gwilym Jenkins. Time series analysis, forecasting and control, 1994.

10. Johan Karlsson, Milena Anguelova, and Mats Jirstrand. An efficient method for structural identifiability analysis of large dynamic systems. *IFAC Proceedings Volumes*, 45(16):941–946, 2012.

11. Thomas BL Kirkwood. Evolution of ageing. *Nature*, 270:301–304, 1977.

12. Armin Rashidi, Thomas BL Kirkwood, and Daryl P Shanley. Evolution of asymmetric damage segregation: A modelling approach. *Aging Research in Yeast*, page 315, 2012.

13. JL Snoep, M Mrwebi, JM Schuurmans, JM Rohwer, and MJ Teixeira de Mattos. Control of specific growth rate in saccharomyces cerevisiae. *Microbiology*, 155(5):1699–1707, 2009.
